# Uncovering aquatic toxicity as a specific stressor among multiple environmental drivers affecting the functional structure of French stream macroinvertebrate communities

**DOI:** 10.64898/2026.01.08.698385

**Authors:** Christopher Bosc, Rémi Recoura-Massaquant, Benjamin Alric, Jérémy Piffady, Olivier Geffard, Arnaud Chaumot

## Abstract

1. Since recent years, freshwater ecosystems such as rivers are subjected to increasing pressures from human-induced drivers of changes, including chemical contamination, warming and nutrient enrichment. Among those, toxic chemicals are particularly concerning, as aquatic contamination is strongly increasing but their effects on ecosystem structure and functioning remain poorly understood and difficult to predict. Yet, those are expected to impact freshwater ecological communities, notably through selecting species exhibiting particular biological traits. A growing literature tested predictions on the effects of multiple environmental drivers, including chemicals, on the functional structure of macroinvertebrate communities, but disentangling the respective influences of the different drivers on the observed patterns usually remains challenging.
2. Here, we used multivariate analyses with variation partitioning to estimate the independent and combined effects of toxic contamination, water temperature and nutrient load on the species abundances and biological traits of macroinvertebrate communities in 76 French streams. Toxic contamination was quantified based on the *Gammarus* feeding inhibition indicator recorded during repeated *in situ* biotests. We then compared trait responses with expectations from a priori predictions derived from the literature, and looked at the link between taxon turnover and taxonomic and functional diversity metrics using linear models.
3. Our results suggest that, when considered independently, each driver had specific impact on invertebrate taxa and biological traits, with different consequences on community composition and diversity. Notably, we showed significant changes in taxonomic composition linked to toxicity, associated with the reduction of both taxonomical and functional richness, but no significant effect on biological trait composition. In contrast, temperature and moreover nutrient load strongly effected traits, but with variables associations with diversity metrics depending on driver. Finally, when all three drivers were considered in combination, patterns were similar to those observed for temperature, suggesting dominance rather than additive effects between drivers.
4. Our study reveals particular negative impacts of toxicity on macroinvertebrate communities in contrast with other drivers, which highlights the potential risk posed by contaminants for freshwater biodiversity.

## Introduction

Freshwater ecosystems, such as rivers and streams, are facing a diversity of human-related pressures including climate changes, hydrological changes, land use changes, chemical inputs and aquatic invasive species, which can potentially hamper past and current efforts to preserve aquatic biodiversity (Carpenter, Stanley & Vander Zanden, 2011; Haase *et al*., 2023). Among those, synthetic chemicals are particularly concerning because, while the number of novel chemical compounds is rapidly increasing worldwide, their effects on organisms and ecosystems are still poorly understood (Steffen *et al*., 2015; Bernhardt, Rosi & Gessner, 2017; Persson *et al*., 2022). Still, many studies around the world suggests that toxic chemicals such as pesticides or metals could have adverse effects on the diversity of aquatic communities, notably macroinvertebrates (Roline, 1988; Hickey & Clements, 1998; Liess & von der Ohe, 2005; Masson *et al*., 2010; Schäfer *et al*., 2011; Kuzmanovic *et al*., 2017; Solis *et al*., 2018; Alric, Geffard & Chaumot, 2022). For instance, in Europe, a recent study observed that the recovery of river macroinvertebrates diversity initiated at the 1970s has stalled since 2010 (Haase *et al*., 2023). This was particularly the case in rivers downstream of dams, urban areas and croplands, where chemical pollution and nutrient enrichment is expected, but also in sites experiencing a faster rate of warming. In those areas, some changes in functional diversity (e.g. decreased functional richness but increased functional evenness) may reflect selection for a subset of traits that confer tolerance (Haase *et al*., 2023).

Generally, disturbed and/or variable environments are deemed to drive the evolution of specific adaptations in organisms, with predictable consequences on community structure and ecosystem functioning (Habitat templet; Southwood, 1977; Odum, 1985). Notably, greater proportion of r-strategists (short generation times, high fecundity), high dispersal, small size species and shorter trophic chains are expected in stressed ecosystems. These predictions have later been derived for river ecosystems (Townsend & Hildrew, 1994) and notably tested in several studies linking changes in biological traits of aquatic macroinvertebrates with different environmental drivers including temperature, nutrients and chemicals (Bonada, Dolédec & Statzner, 2007; Archaimbault *et al*., 2010; Mondy & Usseglio-Polatera, 2013; Floury *et al*., 2016; Kuzmanovic *et al*., 2017; Berger *et al*., 2018; Meyer *et al*., 2022; Lourenço *et al*., 2023). Although those studies validated predictions for some traits, they could not disentangle for the confounding influence of the different drivers and other environmental variables, which was especially the case for toxic chemicals (e.g. pesticides confounded with nutrient enrichment). Meta-analyses of experimental studies notably suggest that dominance or interactions between stressors are important in freshwater ecosystems (Jackson *et al*., 2016; Birk *et al*., 2020), so observational/correlative data should be handled carefully to avoid misinterpretations.

Novel approaches in the measurement of water toxicity may help to disentangle the effect of toxic chemicals on ecosystems from those of other drivers such as nutrient enrichment. Lately, a new biomonitoring tool using the sentinel species *Gammarus fossarum* (Crustacea) has been developed to measure toxicity. This tool uses *in situ* feeding bioassays in which standardised organisms are encaged directly on the monitored stations (Coulaud *et al*., 2011; Besse *et al*., 2013). It allows the calculation of a feeding inhibition index accounting for confounding environmental factors, and supplying an indicator of chemical toxicity present in the stream station during the exposure period. This notably allowed to link stream water toxicity to changes in taxonomic diversity or composition of macroinvertebrate communities (Sarkis *et al*., 2023; Bosc *et al*., 2025).

Here, we especially intended to disentangle the respective role of water toxicity and other environmental drivers (i.e. temperature and nutrient loads) in structuring river macroinvertebrate communities. For that, we used a community ecology approach to analyse the independent and combined effects of the drivers on both the taxonomic and functional composition of invertebrate communities and compared the associations between drivers and biological traits with a priori expectations derived from the literature.

## Methods

### Study sites

We leveraged the monitoring networks managed by French regional water agencies under the European Water Framework Directive (WFD; European Commission, 2000) to select 76 stations in southern and western France. This selection is based on data availability on toxicity (feeding inhibition estimates from *G. fossarum* bioassays), water temperature, nutrient levels and aquatic macroinvertebrate abundances for all stations. These sampling stations are located in wadable rivers (i.e., less than 120 cm deep). In addition, as toxicity data has only recently become available, the study period was limited to 2019–2022. These same stations have already been used in a previous study to establish a link between the same toxicity indicator and macroinvertebrate abundances (Bosc *et al*., 2025).

### Estimation of chemical toxicity based on in situ feeding bioassays

The measurement of feeding inhibition in *G. fossarum* is a standardized method (AFNOR, 2023) used as a chemical toxicity bioassay in French water body monitoring programs (Dedourge-Geffard *et al*., 2009; Coulaud *et al*., 2011; Chaumot *et al*., 2015). It consists of in situ bioassays with selected male organisms that are exposed at monitored sites for a fixed duration. The protocol used in our study was described in Bosc *et al*., 2025 and references therein. It allows the calculation of a feeding inhibition index (FI) which is calibrated against reference feeding rates for given environmental conditions, ensuring that FI primarily reflects chemical toxicity rather than other factors (e.g., temperature, conductivity) (Coulaud *et al*., 2011; Chaumot *et al*., 2015). Three sampling campaigns per year were carried out for each station during the 2019–2022 period. We calculated the average FI value for each station over that period (Fig. 1).

**Fig. 1.**
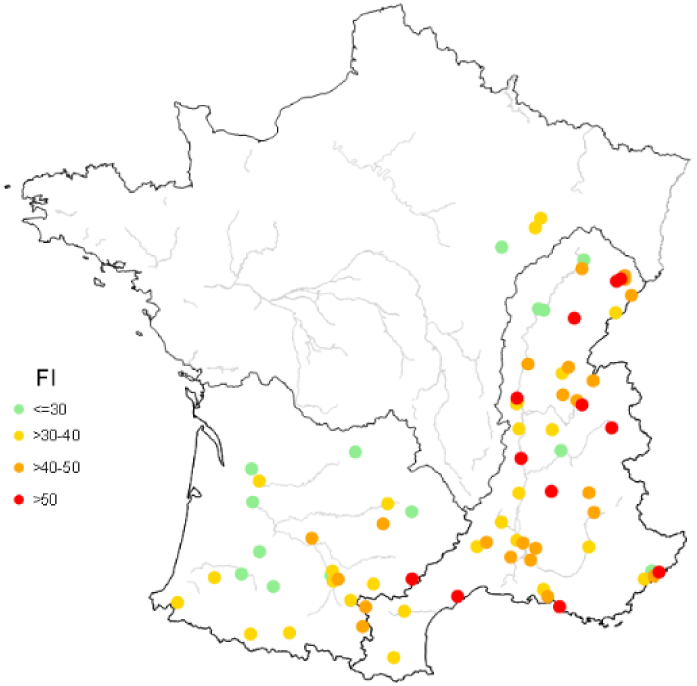
Map of the feeding inhibition index (toxicity in stream water) at the study stations (average 2019-2022). The toxicity level ranges from low toxicity (green) to high toxicity (red).

### Temperature and nutrients

For these 76 stations, we retrieved information on water temperature and nutrient concentration from the French national database “Naïades” (http://www.naiades.eaufrance.fr) for the 2019-2022 period. The nutrients considered in our study were ammonium (NH_4_ ^+^), nitrates (NO_3_ ^-^), nitrites (NO ^-^) and orthophosphates (PO_4_ ^3-^). As with FI index, we considered the average values for each station during the study period. Across all stations, water temperatures were in a range suitable for most taxa (min: 6.9°C; max: 16.6°C), although the highest values may be representative of a thermal stress for temperature-sensitive taxa. The highest nutrient concentration values in our study could be associated with eutrophication (e.g. NH_4_ ^+^ range: 0.01-3.68 mg/L; PO_4_ ^3-^range: 0.02-6.31 mg/L).

### Macroinvertebrate occurrence dataset

From the same national database (Naïades), we extracted aquatic macroinvertebrate abundance data for the same stations and the same period. Sampling was conducted once a year following the IBGN/I2M2 protocol for aquatic macroinvertebrate collection in shallow rivers (AFNOR, 2009). At each station, different habitat fractions were sampled, and invertebrates were stored into three groups based on the habitats they were sampled in (A, B, and C). We focused only on the combined samples from groups B and C, which represent “major habitats”— those sampled based on their representativeness. Only macroinvertebrate taxa identified at the genus and species levels were included in the analysis. When available, species were grouped under their respective genus. Additionally, we identified alien (non-native) taxa using the DAISIE database (Roy *et al*., 2020), considering genus that exclusively contain alien species within the study area. Here too, taxon abundances were averaged in each station in order to obtain a unique station–taxon dataset containing the abundances.

### Macroinvertebrate trait dataset

We obtained a table of the biological traits for the aquatic macroinvertebrate species (taxon–trait category matrix) from the table compiled in Usseglio-Polatera *et al*., 2000 and Tachet *et al*., 2010 which is available on the freshwaterecology.info database (Schmidt-Kloiber & Hering, 2015). For each taxon, affinities to 61 categories of 11 biological traits were coded using a fuzzy-coding technique (Chevenet, Dolédec & Chessel, 1994), where scores ranging from 0 to 5 are assigned to each category, with 0 corresponding to the lowest affinity and 5 to the highest affinity.

### Analyses of the effects of the drivers on the taxonomic and functional compositions of macroinvertebrate communities

Redundancy analyses (RDAs) were performed to quantify the proportion of variance in the taxonomic and functional compositions of aquatic macroinvertebrate communities that were explained by different environmental drivers: toxicity, temperature and nutrients. The explained variance was expressed as adjusted R^2^. Each RDA was based on a response table (taxonomic composition: station–taxon matrix, functional composition: station–trait category matrix) and one, or a combination, of three explanatory variables: the toxicity index (FI), temperature and a composite nutrient variable (hereafter referred as “nutrients”). This nutrient variable corresponds to the scores on the first axis of a principal component analysis (PCA) on the four nutrient variables (NH_4_ ^+^, NO_3_ ^-^, NO_2_ ^-^ and PO_4_ ^3-^), representing 78.3% of the total variance and positively correlated with the four nutrients (Fig. S1). In the case of the response table, a Hellinger transformation was first applied to the abundance data, as recommended to correctly modelling the structure of community data (Legendre & Gallagher, 2001). For the functional composition table, community weighted means (CWM) were calculated for each biological trait category, as the average values of the trait categories for all taxa at each station weighted by relative abundances of taxa. The three explanatory variables were previously transformed in logarithms and normalised using z-scores.

We used a variation partitioning approach to estimate the variance (adjusted R^2^) in taxonomic and functional compositions of macroinvertebrate communities explained independently and conjointly by toxicity, temperature or nutrients. This analysis was performed separately for taxonomic and functional compositions. Variance partitioning is an approach that involves performing several RDAs from which it is possible to estimate the fractions of variances explained independently by each set of predictors and those explained by several sets of predictors in a confounded manner (Borcard *et al*., 1992). Seven RDAs are required to deduce the seven different fractions of variance explained by the three sets of predictors independently and their combinations (see Fig. 2a). We performed permutation tests (n=999) with partial redundancy analyses (pRDAs; Borcard *et al*., 1992) to test the significance of the independent fractions of explained variance in our analyses.

**Fig. 2.**
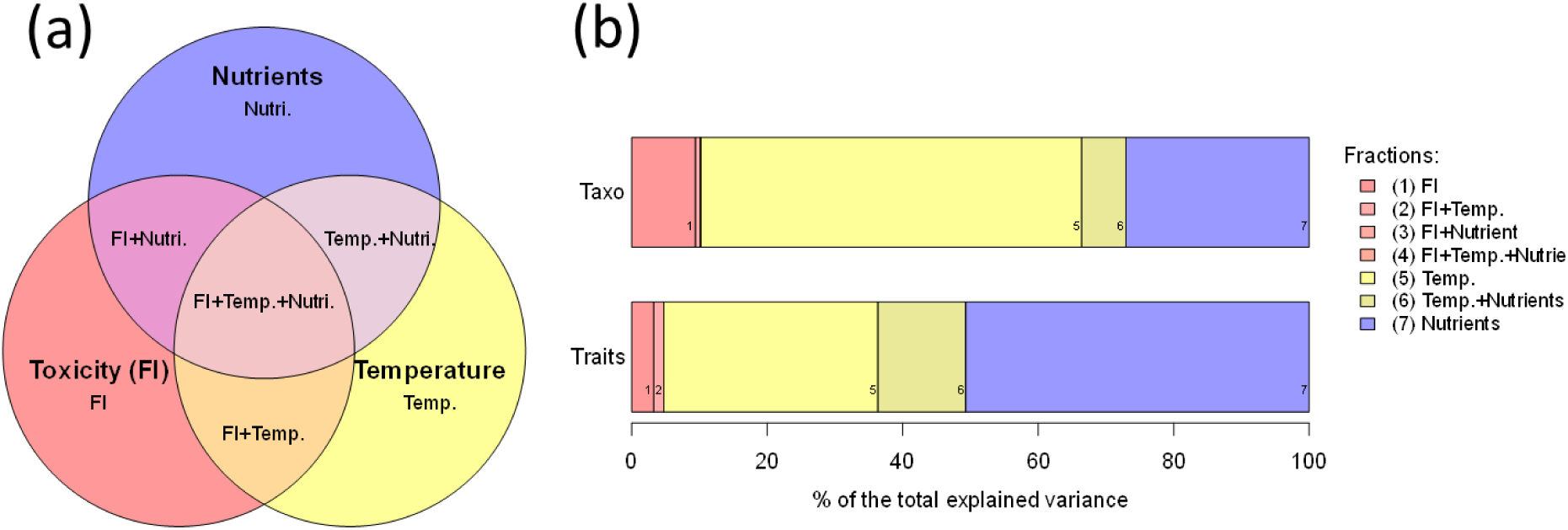
Variation partitioning: (a) concept: each circle represents the variance in macroinvertebrate composition explained by either toxicity, temperature and nutrients, with overlapping parts representing shared fractions of variance. (b) results: variance in taxonomic (top) and functional (bottom) composition explained by the different drivers and representing the different fractions of explained variance. The fractions are obtained from redundancy analyses and are adjusted R-squared values (here expressed as percentages of the total explained variance).

In addition, we estimated the combined effect of the three drivers on the different taxa and trait categories using a composite variable of these factors. A PCA was performed on the three factors, and the second axis, representing 41.2% of the total variance and positively correlated with the three drivers (Fig. S2) (hereafter referred as “combined drivers”), was used as a proxy for their combined effect. For each dataset (taxonomic and functional), an RDA was performed with the composite variable as predictor, and the significance of its effect was tested with permutation tests (n=999). We compared the results of the pRDAs for the drivers considered independently with the results of the RDAs for the drivers in combination by computing Pearson’s correlations of the taxon (or trait category) scores. The aim here was to assess the similarity of the responses of taxa (or trait categories) between the independent and the combined drivers.

### Use of null and a priori models to interpret trait responses to environmental drivers

To help interpreting the relationships between environmental drivers and the composition of biological trait categories in macroinvertebrate communities, we used null models, as well as models of expected trait response to drivers (*a priori* predictions).

*Null models* – It is possible that, under certain conditions, environment–trait relationships are explained essentially by species abundance distribution and not by specific (adaptive) species–trait associations, which is an inherent limitation of RDAs based on CWMs (Lepš & de Bello, 2023). To ensure that the trait responses to environmental drivers revealed by the RDAs are adaptive, we built null models allowing testing the influence of the taxon abundance distribution on the observed environment–trait associations. To do so, several steps were necessary: first, in the taxon–trait category matrix, we randomly reattributed the trait categories to each taxon, while keeping the dependencies between traits (i.e. we did not create new, potentially impossible, combinations of traits). Then, we derived a presence probability matrix from the observed abundance (station–taxon) matrix. This was done by dividing the abundances by the total abundance in the matrix. Based on the given probabilities, we created a new (null) station–taxon matrix by randomly assigning 10000 individuals to stations and taxa. Individuals can be assigned to any station or taxa (i.e. there is no zero probabilities), which allows to obtain a null abundance matrix where abundant taxa have a much greater chance to be found in their original stations than rare taxa. Then, we computed again CWM with the taxon–randomized trait category matrix and the null station–taxon matrix. By doing so, we obtained a station–trait category matrix with similar taxa abundance distribution but randomized traits. Like described above, RDAs were then performed again to see the effects of the different environmental drivers on the randomized trait category composition. The procedure was repeated 1000 times. The comparison of the RDA adjusted R^2^ for the randomized traits with the RDA adjusted R^2^ for the original traits allowed the computation of an overall p-value, while p-values for each trait category were obtained by comparing the randomised scores with the original scores (testing the null hypothesis that dominant taxa explain driver-trait associations). In all the RDAs, positive and negative scores were always associated with respectively low and high driver values.

Standardised trait category scores were then obtained with the formula:

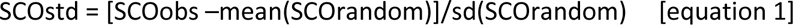

*A priori predictions –* The *a priori* predictions were based on hypotheses about expected trends in stressed or unstable ecosystems (Southwood, 1977; Odum, 1985)(Habitat templet), which were then refined for freshwater communities (Townsend & Hildrew, 1994), but also based on literature related to aquatic macroinvertebrates (Bonada *et al*., 2007; Archaimbault *et al*., 2010; Mondy & Usseglio-Polatera, 2013; Berger *et al*., 2018; Meyer *et al*., 2022). We made predictions for each of the 11 biological traits of invertebrates considered in our study, with predicted responses for 33 trait categories (detailed in Table 1). For these trait categories, we estimated the match of the observed scores with a priori expectations (M) using the standardised scores (SCOstd). For each trait category, if the expected response was positive, then M = SCOstd; if the expected response was negative, M = - SCOstd. Then, for each trait, the median M provided an estimation of the match of the observed scores with a priori expectations (in sd unit). The overall significance of the match for each trait was the median p-value (see above) of its categories.

**Table 1.** A priori predictions of the effect of stress on different biological trait categories of aquatic macroinvertebrates. Derived from references: (1) Odum, 1985; (2) Bonada et al. 2007; (3) Berger et al. 2018; (4) Meyer et al. 2022; (5) Archaimbault et al. 2010; (6) Mondy et Usseglio-Polatera 2013.

| Trait | Category | Stress | Hypothesis | Refs |
| --- | --- | --- | --- | --- |
| Maximal size | < 0.25 cm | + | Stress favors small sizes | 1, 2, 4 |
|  | > 0.25-0.5 cm | + |  | 1, 2, 4 |
|  | > 0.5-1 cm | - |  | 1, 2, 4 |
|  | > 1-2 cm | - |  | 1, 2, 5 |
| Life duration | <= 1 year | + | Stress favors short lifespans | 1 |
|  | > 1 year | - |  | 1 |
| Cycles per year | < 1 | - | Stress favors r-strategies | 1, 2, 4, 5, 6 |
|  | > 1 | + |  | 1, 2, 3, 4, 5, 6 |
| Reproduction | ovoviviparity | + | Egg protection from stress | 2, 3, 4, 5, 6 |
|  | cemented eggs | - |  | 3, 4 |
|  | terrestrial clutches | - |  | 4, 5 |
| Aquatic stages | larva | - | Greater vulnerability of juveniles | 2, 3, 4, 6 |
|  | adult | + |  | 2, 3, 4, 5, 6 |
| Dispersal mode | aquatic passive | + | Selection of optimal locations/avoidance of stressors | 4, 5, 6 |
|  | aquatic active | - |  | 3 |
|  | aerial active | - |  | 3, 4, 5, 6 |
| Resistance forms | eggs | - | Stress favors specialised resistance forms | 3, 4 |
|  | housings | + |  | 2 |
|  | diapause/dormancy | + |  | 2, 4 |
|  | none | - |  | 2, 3 |
| Respiration | gill | + | More efficient when oxygen in water is low | 2, 3, 4 |
|  | spiracle | - |  | 2, 4 |
| Locomotion | crawler | - | Avoidance of pollution in water or dessication | 2, 3, 4, 5 |
|  | burrower | + |  | 3, 4 |
|  | interstitial | + |  | 2, 4 |
| Food | microorganisms | + | Stress shortens food chains/greater sensitivity of predators | 1 |
|  | detritus | + |  | 1 |
|  | macroinvertebrates | - |  | 1 |
|  | vertebrates | - |  | 1 |
| Feeding habits | deposit feeders | + | (as above) | 1 |
|  | filter-feeders | + |  | 1, 3, 4, 5, 6 |
|  | predators | - |  | 1 |
|  | parasites | - |  | 1, 4 |

### Relationship between driver-related changes in taxonomic composition and diversity metrics in macroinvertebrate communities

We selected four diversity metrics yielding information on the taxonomic and functional diversity of the macroinvertebrate communities, which we computed for each station. We considered two independent measures of taxonomic diversity: Taxonomic richness, which was the number of genus, and taxonomic distinctness (Δ*), which is the average taxonomic distance between two individuals of different taxa (Clarke & Warwick, 1998). Δ* was based on a classification table containing the families, orders and classes for each taxon. Likewise, we considered two independent measures of functional diversity: Functional richness (FRic) and functional dispersion (FDis). FRic is a measure of the volume of the multidimensional trait space occupied by all taxa (Villéger, Mason & Mouillot, 2008), and is representative of the number of unique combinations of biological traits found in a community. FDis is the weighted mean distance of individual taxon to the weighted centroid of all taxa within the multidimensional trait space (Laliberté & Legendre, 2010). FDis is thus representative of the distinctness of the different combinations of biological traits found in a community. We examined the relationship between the changes in taxon composition explained by the environmental drivers (i.e. station scores from the RDAs on taxonomic composition) and the four diversity metrics (taxon richness, Δ*, FRic and FDis), by constructing linear models with RDA station scores as predictor variable. Outliers (i.e. scores outside the interquartile range with a factor k=1.5) were not included in the analyses because of their large influence on the results.

pRDAs, RDAs, permutation tests and taxonomic distinctness computation were performed using the vegan package (Oksanen *et al*., 2017) in R software (R Core Team, 2023). We computed CWMs, FRic and FDis with the FD package in R software (Laliberté *et al*., 2014).

## Results

### Independent and confounded influences of the drivers on the taxonomic and functional compositions of macroinvertebrate communities

The results of the variation partitioning procedure (Fig. 2b) show that most of the variance explained by each driver is independent of the effect of the others, this for both the taxonomic and functional compositions of macroinvertebrate communities: the shared (confounded) fractions represent only 7.3% and 14.4% of the total explained variance respectively. With regard to taxonomic composition, pRDAs showed that environmental drivers, either alone or in combination, influence the taxonomic composition of macroinvertebrate communities, with significant effects in both cases (Table S1). The effect of toxicity although significant was reduced compared to other drivers, with temperature having the greatest influence on taxonomic composition. The percentages of the total explained variance attributable to toxicity, temperature and nutrients were respectively of 10.2%, 63.4% and 33.6% (including shared fractions). Concerning the functional composition of macroinvertebrate communities, the results showed that toxicity alone does not significantly explain it(F=1.41; p = 0.156; Table S1), while other drivers and drivers in combination are significant (Table S1). Nutrients had the largest influence on the functional composition, with a relative contribution of 63.6%, followed by temperature (46%) and toxicity (4.8%).

### Responses of macroinvertebrate taxa to environmental drivers

Fig. 3 shows the responses of the different taxa to the different drivers, considered alone (pRDA scores) or in combination (RDA scores) (Details of taxon scores in Table S2). We compared main macroinvertebrate groups (insect orders and other macroinvertebrate classes), as well as native and alien taxa. When we consider the independent association with toxicity, we can see a tendency for the majority of taxonomical groups to be mostly negatively associated with toxicity (indicating sensitivity), but not Plecoptera and Malascostraca that responded mostly positively (indicating tolerance). For temperature, Diptera and more importantly Plecoptera are significantly associated with relatively cold temperatures. For nutrients, Trichoptera, Ephemeroptera and Plecoptera have a significantly negative association with them, while non-insect taxa mostly have a positive (though not significant) association. Finally, when drivers are considered in combination, most insect groups and Clitellata showed a significant negative association with them, while non-insect taxa tended to respond positively (though not significantly). We observed, for all environmental drivers, taken individually or in combination, a tendency for native taxa to be significantly and negatively associated with drivers, whereas a positive (sometimes significant) association was observed for alien taxa.

**Fig. 3.**
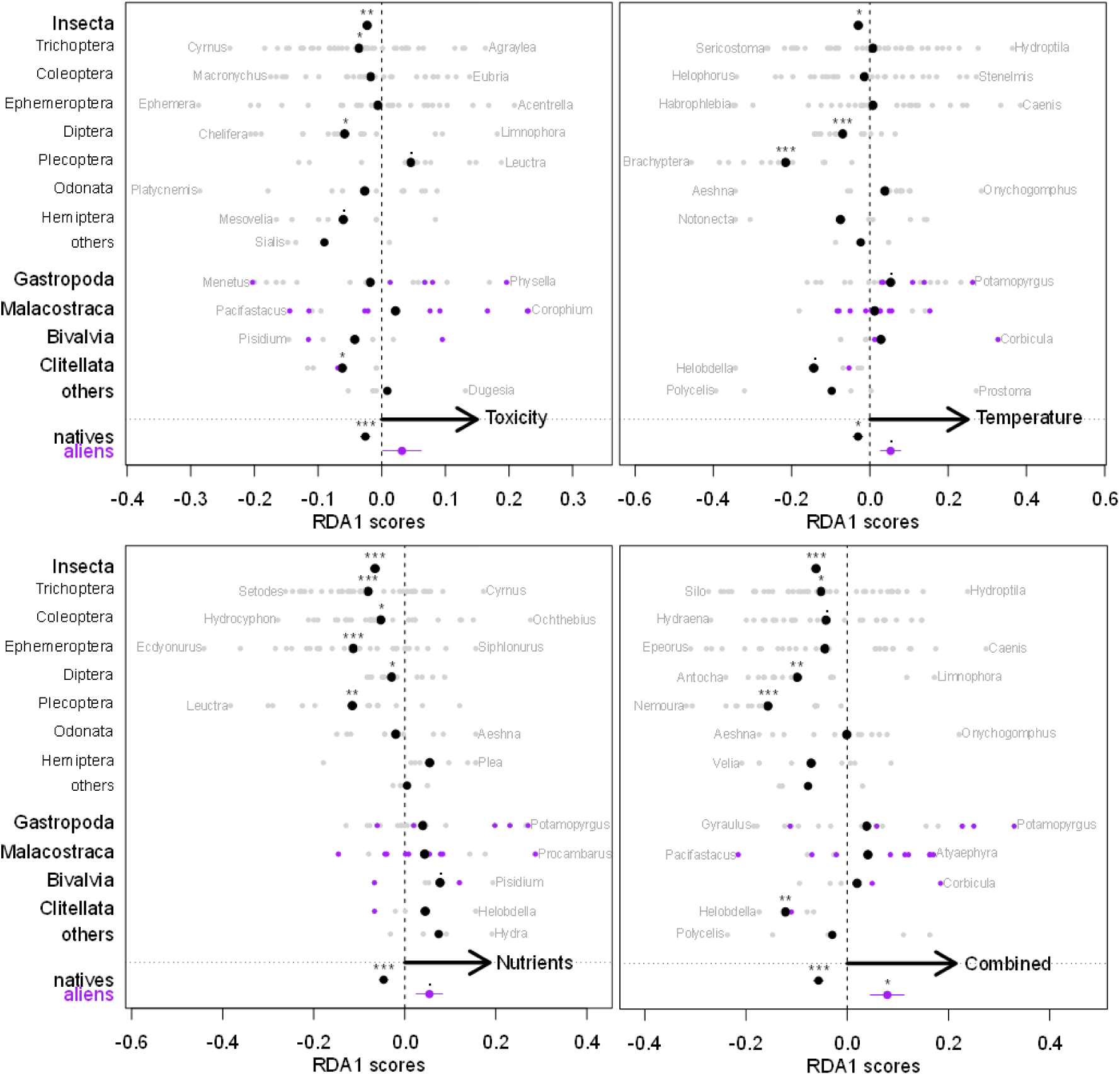
Taxon scores from (partial) redundancy analyses for the different drivers and their combination. The scores are sorted by taxonomic group or origin (alien vs native). Native taxa are figures in gray and alien taxa are figured in purple. Large black points are mean group scores. Significance of group mean deviation from zero : . = p<0,1; * = p<0.05; ** = p<0.01; *** = p<0.001.

Specifically, *Physella* (Gastropoda) and the amphipods *Dikerogammarus* and *Corophium* (Malacostraca) are the alien taxa the most positively associated with toxicity. *Potamopyrgus* (Gastropoda) is positively associated with both temperature and nutrients. *Corbicula* (Bivalvia) is the alien taxa the most positively associated with temperature and the crayfish *Procambarus* (Malacostraca) the most positively associated with nutrients. Overall, the results for combined drivers were highly correlated with those found for temperature (Pearson’s correlation of taxa scores: 0.80; p<0.001), moderately correlated with those for toxicity (0.50; p<0.001) and weakly correlated with those for nutrients (0.17; p=0.018).

### Macroinvertebrate trait responses to environmental drivers compared to null models and a priori predictions

For temperature, nutrients and drivers in combination, we found that observed adjusted R2 values of pRDAs on traits are significantly different from those obtained from null models (Table 2), which confirms the observed associations between those drivers and trait composition, independently of a potential bias due to species abundance distribution.

**Table 2.**
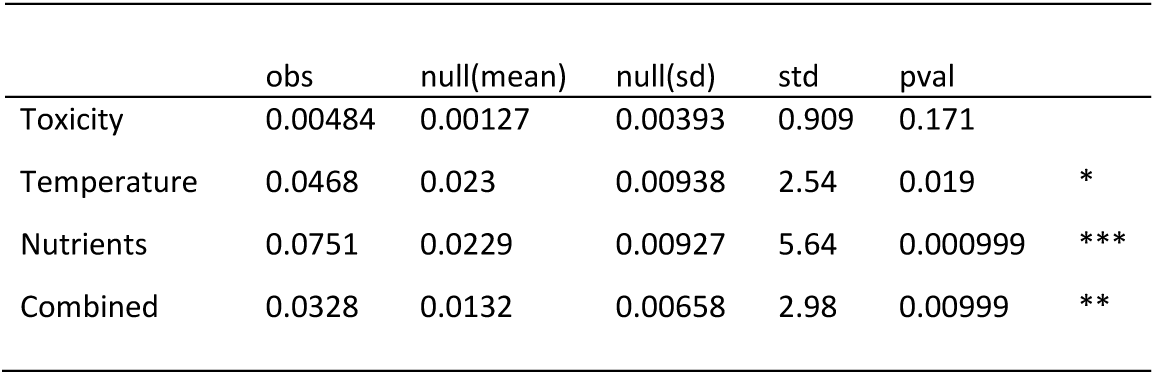
Adjusted R2 values of (p)RDA on biological traits of invertebrates (observed vs null models)

We examined the responses of the different trait categories to environmental drivers (standardised scores SCOstd of pRDAs; Fig. 4), comparing them to *a priori* predictions. Concerning the independent effect of toxicity, we found that most traits tend to fit expectations (8 out of 11), but never significantly (Fig. 4; Table S3). For temperature, all traits tend to fit expectations, but only significantly so for cycles per year with multivoltine taxa associated with elevated temperatures, and locomotion with interstitial taxa associated with low temperatures (Fig. 4; Table S3). Maximal size, respiration and food are traits with a marginally significant (p<=0.1) association with temperature.

**Fig. 4.**
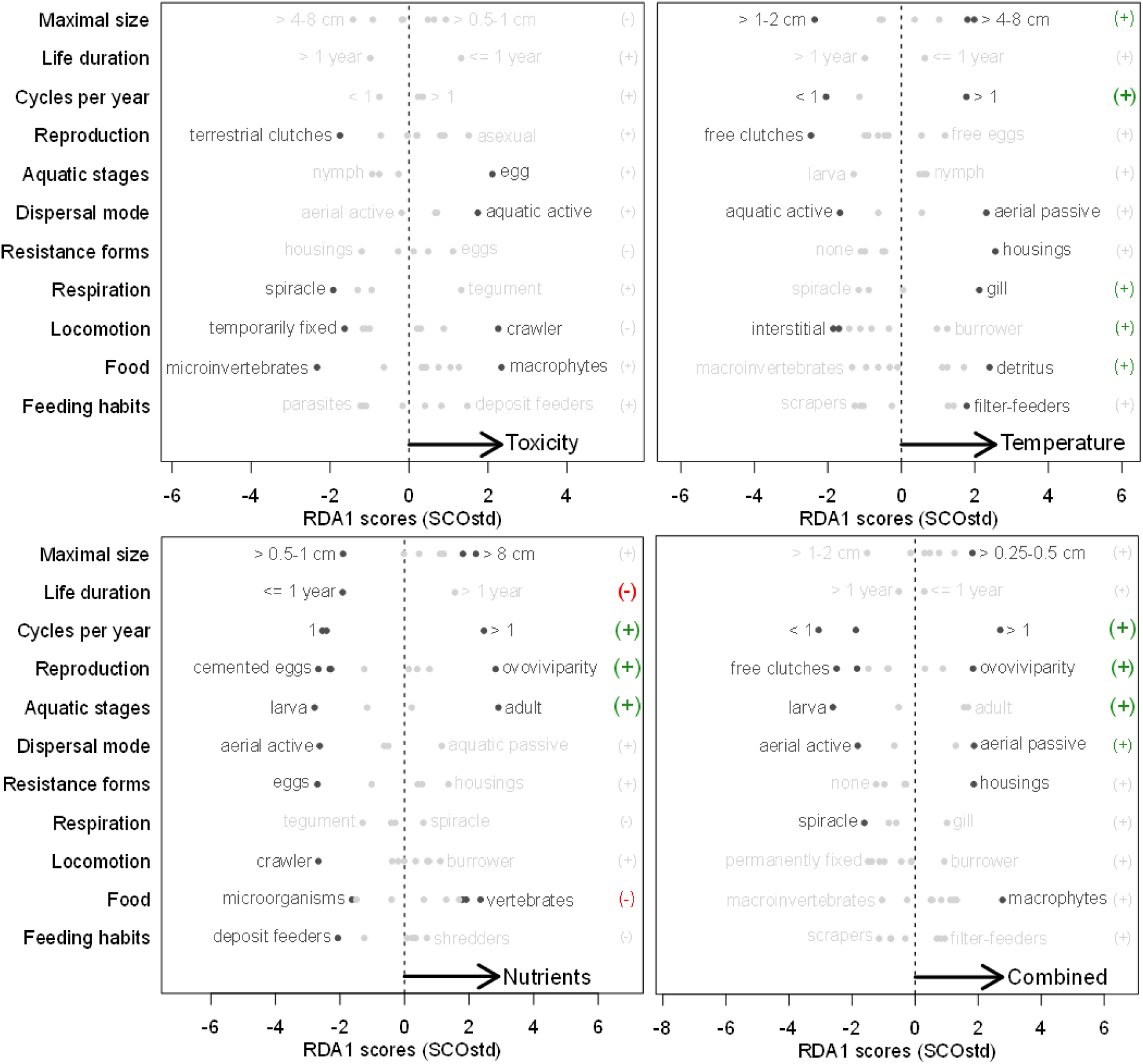
Trait category scores from (partial) redundancy analyses for the different drivers and their combination. Black points are trait categories significantly different from null models. The green (+) represent trait responses matching a priori predictions (with p<=0.1; see Table 1), while red (-) are traits responding in the opposite way. Greys (+) and (-) are traits not significantly matching predictions (with p>0,1). The sizes of the (+) and (-) are also function of the significance (the larger the most significant).

For nutrients, 7 out of 11 traits fit expectations. Especially, cycles per year, reproduction, and aquatic stages respond strongly and significantly to nutrient gradient and accordingly to expectations (Fig. 4; Table S3), while life duration and food have (marginally) significant associations with nutrients but opposite to predicted. Finally, trait responses to combined driver tend to fit expectation for all traits, but most strongly and significantly so for cycles per year, reproduction and aquatic stages (Fig. 4; Table S3), and marginally significantly so for dispersal. These results for combined drivers are highly correlated with those found for temperature (Pearson’s correlations of trait modality scores: 0.80; p<0.001), well correlated with those for nutrients (0.60; p<0.001) but uncorrelated with those for toxicity (0.19; p=0.143).

### Association between driver-related taxon turnover and diversity metrics in macroinvertebrate communities

Changes in taxonomic composition of macroinvertebrate communities related to toxicity and nutrients (pRDA station scores) are associated with significant reductions in taxonomic richness (number of Genus) (Fig. 5). However, nutrients-related changes in composition are also associated with a sharp increase in taxonomic distinctness (Δ*), while Δ* did not significantly change with toxicity-related changes in composition. The changes in taxonomic composition related to temperature and drivers in combination are not associated with significant changes in taxonomic richness or distinctness (Fig. 5). Then, the changes in taxonomic composition related to toxicity are also associated with significant reductions in both functional richness (FRic) and dispersion (FDis), while FRic and FDis increase with temperature-related changes in composition. Finally, the changes in taxonomic composition related to nutrients and drivers in combination are associated with increased functional dispersion, but no significant changes in functional richness (Fig. 5).

**Fig. 5.**
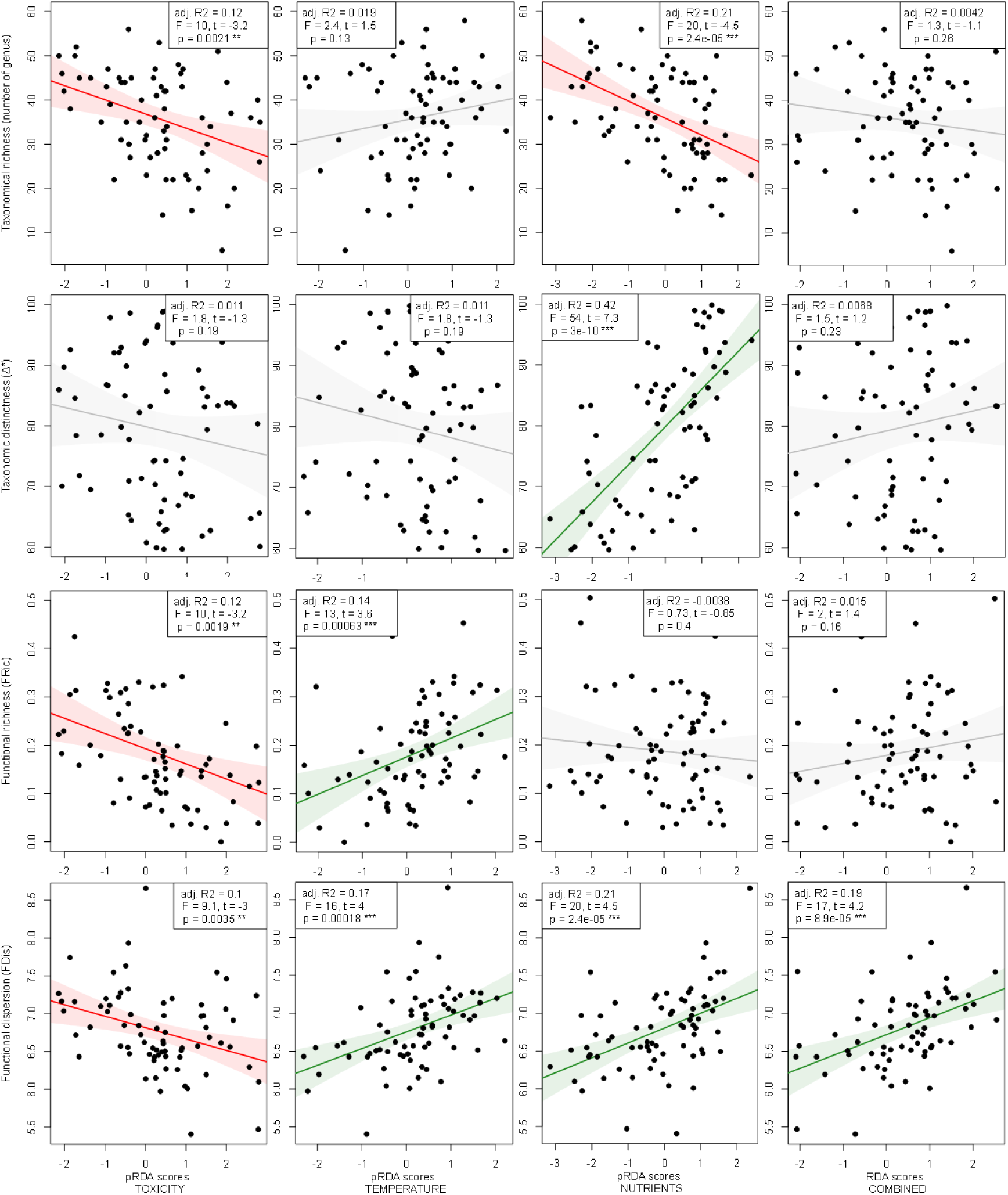
Relationship between station scores (taxonomic tunover linked to the different drivers and their combination) and diversity metrics (taxonomic richness and distinctness; functional richness and dispersion). Linear regression lines with 95% confidence intervals.

## Discussion

In this study, we estimated the relative importance of toxicity, temperature and nutrients in explaining changes in the taxonomic and functional compositions of aquatic macroinvertebrate communities using RDAs, and tested the association between those changes and diversity metrics using linear models. Our variation partitioning procedure allowed us to identify the associations between macroinvertebrate community compositions and the drivers, independently or in combination. Our results suggest that each driver had specific impact on macroinvertebrate taxa and biological traits, with different consequences on community composition and diversity metrics.

Notably, we showed significant changes in taxonomic composition linked to toxicity, and showed that these changes were associated with a reduction of both taxonomic and functional diversity metrics, but we did not detect significant effects of toxicity on functional composition. Temperature was associated with the greatest changes in taxonomic composition, and with increased functional diversity (richness and dispersion), whereas nutrients were the main contributor to changes in functional composition, associated with increased functional divergence, but with reduced taxonomic diversity. The combined variables significantly changed both taxonomic and functional compositions, with a positive effect on functional divergence but none on other diversity metrics.

Our results concerning toxicity–macroinvertebrate community associations suggest chemical contamination triggers specific taxonomic changes (significant changes in taxonomic composition), with more taxa loss than gain (declining taxonomic richness), but seemingly weak selection of the biological traits (no effects of toxicity on functional composition) with yet more trait loss than gain (declining functional richness). These results are in accordance with preliminary work conducted at a smaller scale suggesting that metals affect more the taxonomic structure of stream macroinvertebrate communities than they affect their biological traits (Alric *et al*., 2022). However, this does not mean that the responses of macroinvertebrates to toxicity is never selective or adaptive, merely that it does not imply the traits considered in our study. Other biological traits that are more closely linked to the internal physiology of invertebrates would likely respond to toxic chemicals, but these are usually not included in classical trait databases and require substantial effort to measure. For instance, Buchwalter & Luoma (2005) showed that general traits such as body size or gill size did not influence dissolved Cadmium and Zinc uptakes, whereas specific traits associated with osmoregulatory mechanisms such as the number of ion channels were important. Another study showed that sensitivity to pesticides was well correlated with traits related to somatic maintenance (Baas & Kooijman, 2015), while Ippolito, Todeschini & Vighi (2012) suggest behavioural complexity (as a proxy of nervous system complexity) could respond to toxicity. In addition, traits such as maximal body size could be replaced by more sensitive traits such as actual body size, which would more closely respond to variation of toxicity (Wiberg-Larsen *et al*., 2016). Also, it must be noted that some life-history traits may differ between populations within the same species because of selection by contaminants (e.g. reduced body size along a metal contamination gradient; Lalouette *et al*., 2024). In that case, the average value of the trait observed at the species (and higher taxonomic) level might not be relevant for assessing the effect of toxicity.

When comparing the results of the RDAs on taxonomical vs functional compositions of macroinvertebrate communities, we saw that temperature was associated with greater taxon-related than trait-related changes, even though trait changes mostly followed expected trends and taxon-related changes were positively associated with functional diversity metrics. Overall, these results confirm previous findings in the literature, notably the association of high temperatures with multivoltinism, shorter trophic length and higher diversity (Bonada *et al*., 2007; Bonacina *et al*., 2023). This confirms temperature induces adaptive response of macroinvertebrates, which was expected since temperature represents a natural environmental gradient acting at an evolutionary timescale.

Contrary to the results observed for temperature, nutrients were associated with greater changes in functional than taxonomical compositions, which was accompanied by positive associations of taxon-related changes with taxonomic distinctness and functional dispersion, but negative association with taxonomic richness. Some traits followed trends expected in stressed ecosystems (e.g. multivoltinism and ovoviviparity), but others followed opposite trends (e.g. feeding habits). In our study, we observed nutrient–macroinvertebrate community associations comparable to those observed in the literature (e.g. Weijters *et al*., 2009; Mondy & Usseglio-Polatera, 2013; Floury *et al*., 2016; Berger *et al*., 2018), notably the negative association of nutrients with taxonomic richness, with particular sensitivity of the Ephemeroptera, Trichoptera and Plecoptera (EPT taxa), and positive association with multivoltinism and ovoviviparity. However, our study highlighted some aspects of the effects of nutrients on macroinvertebrate communities not necessarily seen previously. Notably, our results suggest that nutrients could reduce species richness while improving functional diversity (i.e. dispersion) with the RDAs suggesting strong changes in trait composition within the community. This means that nutrients act as an important selective agent on macroinvertebrates, with contrasting effects on their taxonomic and functional diversity.

In addition, our results showed that the effects of the three drivers in combination were mainly comparable to the independent effect of temperature, this for both the taxonomic and functional compositions of macroinvertebrate communities, and despite nutrients having the strongest effect on the functional composition when considered alone. One explanation could be that warm-adapted taxa possess combination of traits that makes them better able to cope with other drivers of stress (e.g. multivoltine taxa are favoured in high temperature and high nutrients/eutrophic condition), so that the dominance of these taxa in warm waters would diminish the effects of other drivers. This would suggest that the net effect of temperature and other drivers is mainly antagonistic (i.e. less than the sum of their expected combined effects, with the dominance of temperature). This contrasts the results of previous meta-analyses of the effect of warming and other stressors that found either additive net effect of warming and nutrient enrichment (Jackson *et al*., 2016) or dominance of local stressors over warming (Morris *et al*., 2022). However, discrepancies between previous studies and our results could be linked to differences in the response variables considered (e.g. single species vs community structure), type of study (experimental vs observational) or scale (local/temporal vs regional/spatial). Also, overall, studies on the interactive effect of temperature/warming and chemical pollution are still lacking (Polazzo *et al*., 2022).

In our study, one notable commonality of the effects of drivers on invertebrates is that they all overall favoured alien taxa over native ones. This was found for toxicity, temperature and nutrients independently and in combination. Those results are in line with previous findings suggesting warming, eutrophication, chemical pollution or changes in salinity can favour alien taxa (Grabowski, Bacela & Konopacka, 2007; Paillex *et al*., 2017; Collas *et al*., 2018; Sarà *et al*., 2018; Cieplok, Spyra & Czerniawski, 2023).

## Conclusion

Results from our study suggest that toxicity has contrasting negative effects on macroinvertebrate diversity compared to temperature and nutrients, even though toxicity contribution to changes in macroinvertebrate communities is low and not linked to particular biological traits. This implies that the effects of toxic contamination on ecosystems may be particularly hard to predict, especially when other widespread drivers of changes such as warming or nutrient enrichment co-occur. Toxic contamination of rivers was flagged as a plausible cause for some of the unexplained low recovery rate of macroinvertebrate communities in European human-impacted areas (Haase *et al*., 2023). Our results point to particular negative impacts of toxicity which supports that explanation and advocate for a better consideration of the impact of contaminants on freshwater biodiversity in environmental policies and conservation plans.

## Supporting information

R scripts

## Acknowledgments

We are thankful to RMC water agency for funding, Biomae for simplified access to data and metadata, as well as Laurent Valette and Martial Ferréol for additional information about data.

## Author contributions

**Christopher Bosc**: Conceptualization, Methodology, Formal analysis, Writing - Original Draft. **Rémi Recoura-Massaquant**: Data Curation, Writing - Review & Editing. **Benjamin Alric**: Conceptualization, Methodology, Writing - Review & Editing. **Jérémy Piffady**: Conceptualization, Methodology, Writing - Review & Editing. **Olivier Geffard**: Conceptualization, Methodology, Supervision, Writing - Review & Editing. **Arnaud Chaumot**: Conceptualization, Methodology, Supervision, Writing - Review & Editing.

