## Supplementary figures and tables for "Uncovering aquatic toxicity as a specific stressor among multiple environmental drivers affecting the functional structure of French stream macroinvertebrate communities"

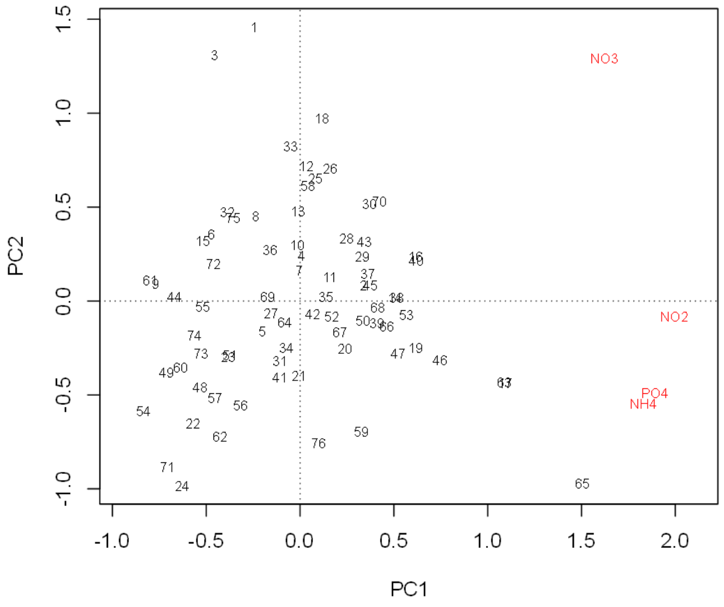


Fig. S1. Results of a principal component analysis on four nutrient variables: ammonium (NH4), nitrates (NO3), nitrites (NO2) and orthophosphates (PO4).

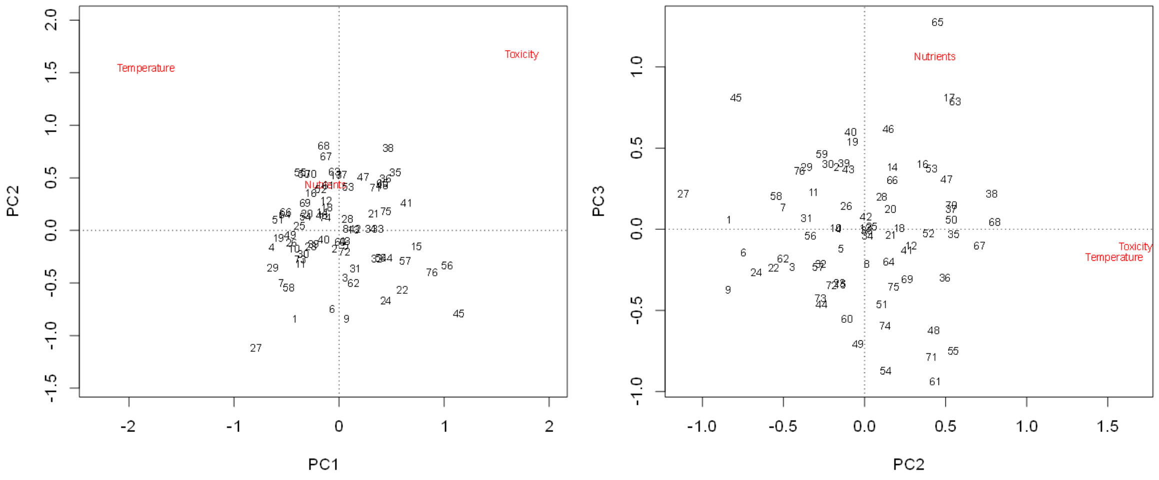


Fig. S2. Results of a principal component analysis on the three environmental drivers: Toxicity, Nutrients and Temperature.

**SUPPLEMENTARY TABLES**

Table S1. Significance of (p)RDAs based on permutation tests.

| Response | Driver | Df | Variance | F | Pr(>F) | |
|---|---|---|---|---|---|---|
| Taxonomic | Toxicity | 1 | 3.53422414932085 | 1.4285971995774 | 0.021 | * |
| | Temperature | 1 | 8.80667406972342 | 3.55981663359281 | 0.001 | *** |
| | Nutrients | 1 | 5.51877938777919 | 2.23079025137154 | 0.001 | *** |
| | Combined | 1 | 6.24392974826154 | 2.42220759088605 | 0.001 | *** |
| Functional | Toxicity | 1 | 0.982750781862997 | 1.41311974733316 | 0.156 | |
| | Temperature | 1 | 3.47636909867427 | 4.99875035768758 | 0.001 | *** |
| | Nutrients | 1 | 5.15720042144077 | 7.41565602489528 | 0.001 | *** |
| | Combined | 1 | 2.78909540233441 | 3.54560818457068 | 0.001 | *** |

Table S2. Taxon scores from (p)RDAs.

| | Toxicity | Temperature | Nutrients | Combined | group | origin |
|---|---|---|---|---|---|---|
| Brachyptera | 0.056 | -0.457 | 0.121 | -0.218 | Plecoptera | native |
| Rhabdiopteryx | 0.053 | -0.116 | -0.059 | -0.063 | Plecoptera | native |
| Taeniopteryx | 0.037 | -0.046 | -0.019 | -0.012 | Plecoptera | native |
| Amphinemura | 0.039 | -0.323 | -0.081 | -0.22 | Plecoptera | native |
| Nemoura | -0.13 | -0.361 | 0.039 | -0.318 | Plecoptera | native |
| Protonemura | 0.148 | -0.276 | -0.225 | -0.168 | Plecoptera | native |
| Leuctra | 0.188 | -0.234 | -0.383 | -0.174 | Plecoptera | native |
| Capnia | 0.053 | -0.116 | -0.059 | -0.063 | Plecoptera | native |
| Capnioneura | 0.053 | -0.116 | -0.059 | -0.063 | Plecoptera | native |
| Isoperla | 0.068 | -0.385 | -0.078 | -0.239 | Plecoptera | native |
| Perlodes | 0.138 | -0.119 | -0.197 | -0.06 | Plecoptera | native |
| Dinocras | 0.077 | -0.197 | -0.3 | -0.195 | Plecoptera | native |
| Perla | -0.032 | -0.254 | -0.29 | -0.306 | Plecoptera | native |
| Chloroperla | 0.053 | -0.116 | -0.059 | -0.063 | Plecoptera | native |
| Siphonoperla | -0.114 | -0.116 | -0.077 | -0.188 | Plecoptera | native |
| Rhyacophila lato-sensu | 0.127 | -0.261 | -0.246 | -0.181 | Trichoptera | native |
| Glossosoma | -0.099 | -0.166 | -0.121 | -0.228 | Trichoptera | native |
| Agapetus | -0.123 | -0.099 | -0.206 | -0.235 | Trichoptera | native |
| Orthotrichia | -0.111 | 0.133 | -0.112 | -0.034 | Trichoptera | native |
| Ithytrichia | -0.027 | 0.098 | -0.026 | 0.036 | Trichoptera | native |
| Oxyethira | 0.042 | 0.106 | 0.044 | 0.118 | Trichoptera | native |
| Hydroptila | 0.058 | 0.364 | -0.117 | 0.238 | Trichoptera | native |
| Agraylea | 0.163 | 0.186 | -0.227 | 0.15 | Trichoptera | native |
| Allotrichia | -0.074 | 0.047 | -0.097 | -0.06 | Trichoptera | native |
| Chimarra | -0.011 | 0.277 | -0.231 | 0.086 | Trichoptera | native |
| Philopotamus | -0.024 | -0.019 | -0.22 | -0.116 | Trichoptera | native |
| Wormaldia | -0.062 | 0.066 | -0.138 | -0.054 | Trichoptera | native |
| Hydropsyche | -0.025 | 0.228 | -0.138 | 0.08 | Trichoptera | native |

| | | | | | | |
|---|---|---|---|---|---|---|
| Cheumatopsyche | -0.023 | 0.149 | 0.017 | 0.09 | Trichoptera | native |
| Cyrnus | -0.239 | 0.023 | 0.173 | -0.085 | Trichoptera | native |
| Polycentropus | -0.015 | 0.172 | -0.182 | 0.032 | Trichoptera | native |
| Neureclipsis | -0.028 | -0.087 | 0.004 | -0.076 | Trichoptera | native |
| Psychomyia | 0.001 | -0.008 | 0.057 | 0.018 | Trichoptera | native |
| Lype | -0.184 | 0.038 | 0.021 | -0.096 | Trichoptera | native |
| Tinodes | 0.065 | -0.185 | 0.084 | -0.045 | Trichoptera | native |
| Metalype | -0.082 | -0.033 | -0.094 | -0.117 | Trichoptera | native |
| Ecnomus | 0.02 | 0.106 | 0.042 | 0.101 | Trichoptera | native |
| Brachycentrus | -0.021 | -0.086 | -0.087 | -0.107 | Trichoptera | native |
| Micrasema | 0.039 | -0.201 | -0.125 | -0.156 | Trichoptera | native |
| Goera | -0.082 | 0.025 | 0.051 | -0.022 | Trichoptera | native |
| Silo | -0.15 | -0.195 | -0.097 | -0.274 | Trichoptera | native |
| Lepidostoma | -0.146 | -0.218 | -0.004 | -0.25 | Trichoptera | native |
| Lasiocephala | 0.037 | -0.188 | -0.122 | -0.147 | Trichoptera | native |
| Athripsodes | -0.054 | 0.079 | -0.145 | -0.043 | Trichoptera | native |
| Mystacides | -0.127 | 0.202 | 0.014 | 0.051 | Trichoptera | native |
| Ceraclea | 0.115 | 0.1 | -0.123 | 0.1 | Trichoptera | native |
| Triaenodes | 0.023 | 0.081 | -0.011 | 0.066 | Trichoptera | native |
| Oecetis | -0.164 | 0.022 | 0.023 | -0.092 | Trichoptera | native |
| Setodes | 0.13 | 0.148 | -0.262 | 0.088 | Trichoptera | native |
| Sericostoma | 0.032 | -0.265 | -0.227 | -0.244 | Trichoptera | native |
| Beraea | -0.108 | -0.105 | -0.098 | -0.185 | Trichoptera | native |
| Beraeodes | -0.041 | -0.08 | 0.007 | -0.08 | Trichoptera | native |
| Beraeamyia | -0.074 | 0.047 | -0.097 | -0.06 | Trichoptera | native |
| Odontocerum | -0.068 | -0.108 | -0.162 | -0.184 | Trichoptera | native |
| Siphlonurus | -0.008 | -0.344 | 0.156 | -0.174 | Ephemeroptera | native |
| Baetis | 0.072 | -0.075 | -0.26 | -0.102 | Ephemeroptera | native |
| Centroptilum | -0.206 | -0.038 | 0.003 | -0.17 | Ephemeroptera | native |
| Cloeon | 0.091 | 0.16 | 0.01 | 0.175 | Ephemeroptera | native |
| Procloeon | -0.193 | 0.248 | -0.187 | -0.045 | Ephemeroptera | native |
| Oligoneuriella | 0.168 | 0.124 | -0.284 | 0.09 | Ephemeroptera | native |
| Epeorus | 0.01 | -0.347 | -0.215 | -0.309 | Ephemeroptera | native |
| Rhithrogena | 0.027 | -0.299 | -0.249 | -0.278 | Ephemeroptera | native |
| Ecdyonurus | 0.143 | -0.015 | -0.441 | -0.083 | Ephemeroptera | native |
| Heptagenia | 0.018 | 0.206 | -0.061 | 0.126 | Ephemeroptera | native |
| Ephemerella | 0.017 | -0.098 | -0.188 | -0.128 | Ephemeroptera | native |
| Caenis | 0.067 | 0.385 | -0.076 | 0.274 | Ephemeroptera | native |
| Brachycercus | -0.016 | -0.022 | -0.026 | -0.037 | Ephemeroptera | native |
| Choroterpes | -0.037 | 0.334 | -0.361 | 0.055 | Ephemeroptera | native |
| Paraleptophlebia | -0.157 | -0.021 | 0.027 | -0.114 | Ephemeroptera | native |
| Habroleptoides | -0.064 | -0.157 | -0.133 | -0.203 | Ephemeroptera | native |
| Habrophlebia | -0.039 | -0.35 | -0.017 | -0.268 | Ephemeroptera | native |
| Ephoron | -0.061 | 0.237 | -0.099 | 0.076 | Ephemeroptera | native |
| Ephemera | -0.288 | -0.054 | -0.019 | -0.247 | Ephemeroptera | native |
| Potamanthus | -0.032 | 0.117 | 0.091 | 0.091 | Ephemeroptera | native |

| | | | | | | |
|---|---|---|---|---|---|---|
| Gyrinus | -0.015 | -0.227 | 0.151 | -0.103 | Coleoptera | native |
| Orectochilus | -0.099 | -0.082 | -0.153 | -0.185 | Coleoptera | native |
| Haliplus | 0.055 | -0.241 | 0.124 | -0.073 | Coleoptera | native |
| Peltodytes | 0.081 | 0.229 | -0.212 | 0.127 | Coleoptera | native |
| Brychius | 0.104 | -0.113 | -0.117 | -0.048 | Coleoptera | native |
| Oreodytes | -0.032 | -0.142 | -0.073 | -0.146 | Coleoptera | native |
| Laccophilus | 0.003 | 0.151 | -0.079 | 0.072 | Coleoptera | native |
| Platambus | -0.056 | -0.121 | -0.063 | -0.145 | Coleoptera | native |
| Laccobius | -0.166 | -0.003 | 0.023 | -0.11 | Coleoptera | native |
| Helophorus | -0.028 | -0.341 | 0.122 | -0.199 | Coleoptera | native |
| Hydraena | -0.156 | -0.197 | -0.07 | -0.27 | Coleoptera | native |
| Ochthebius | 0.069 | -0.09 | 0.276 | 0.097 | Coleoptera | native |
| Helichus | -0.044 | 0.136 | -0.073 | 0.03 | Coleoptera | native |
| Dryops | 0.081 | 0.247 | -0.191 | 0.147 | Coleoptera | native |
| Potamophilus | -0.119 | 0.108 | 0.016 | -0.006 | Coleoptera | native |
| Stenelmis | -0.053 | 0.272 | 0.009 | 0.147 | Coleoptera | native |
| Elmis | 0.016 | -0.152 | -0.13 | -0.141 | Coleoptera | native |
| Esolus | -0.15 | -0.002 | -0.147 | -0.166 | Coleoptera | native |
| Dupophilus | -0.035 | -0.107 | -0.075 | -0.125 | Coleoptera | native |
| Oulimnius | -0.01 | 0.005 | 0.073 | 0.025 | Coleoptera | native |
| Limnius | -0.153 | 0.037 | -0.2 | -0.162 | Coleoptera | native |
| Normandia | 0.117 | 0.039 | -0.128 | 0.058 | Coleoptera | native |
| Riolus | -0.025 | -0.095 | -0.066 | -0.107 | Coleoptera | native |
| Macronychus | -0.175 | 0.084 | 0.023 | -0.059 | Coleoptera | native |
| Eubria | 0.138 | -0.045 | -0.026 | 0.057 | Coleoptera | native |
| Hydrocyphon | 0.068 | 0.02 | -0.279 | -0.049 | Coleoptera | native |
| Calopteryx | -0.179 | 0.103 | 0.085 | -0.025 | Odonata | native |
| Platycnemis | -0.286 | 0.078 | 0.063 | -0.125 | Odonata | native |
| Boyeria | 0.087 | 0.051 | -0.119 | 0.049 | Odonata | native |
| Aeshna | -0.008 | -0.344 | 0.156 | -0.174 | Odonata | native |
| Gomphus | -0.062 | 0.101 | -0.003 | 0.022 | Odonata | native |
| Ophiogomphus | 0.034 | 0.068 | -0.016 | 0.063 | Odonata | native |
| Onychogomphus | 0.068 | 0.285 | -0.044 | 0.221 | Odonata | native |
| Cordulegaster | -0.078 | -0.05 | -0.149 | -0.147 | Odonata | native |
| Oxygastra | 0.034 | 0.068 | -0.016 | 0.063 | Odonata | native |
| Somatochlora | -0.027 | -0.057 | -0.023 | -0.066 | Odonata | native |
| Orthetrum | 0.032 | 0.083 | -0.127 | 0.028 | Odonata | native |
| Sialis | -0.148 | -0.03 | -0.01 | -0.128 | Megaloptera | native |
| Micronecta | -0.141 | 0.139 | 0.033 | 0.006 | Hemiptera | native |
| Aphelocheirus | -0.084 | 0.103 | 0.014 | 0.015 | Hemiptera | native |
| Notonecta | -0.008 | -0.344 | 0.156 | -0.174 | Hemiptera | native |
| Plea | -0.008 | -0.344 | 0.156 | -0.174 | Hemiptera | native |
| Gerris | -0.099 | 0.007 | 0.138 | -0.011 | Hemiptera | native |
| Hydrometra | 0.084 | 0.146 | -0.179 | 0.086 | Hemiptera | native |
| Mesovelia | -0.166 | -0.003 | 0.023 | -0.11 | Hemiptera | native |
| Velia | -0.06 | -0.306 | 0.097 | -0.209 | Hemiptera | native |

| | | | | | | |
|---|---|---|---|---|---|---|
| Blephariceera | -0.028 | -0.138 | -0.081 | -0.143 | Diptera | native |
| Liponeura | -0.032 | -0.142 | -0.073 | -0.146 | Diptera | native |
| Antocha | -0.189 | -0.13 | -0.05 | -0.24 | Diptera | native |
| Dicranota | -0.07 | -0.063 | -0.049 | -0.111 | Diptera | native |
| Eloeophila | -0.124 | -0.088 | -0.046 | -0.164 | Diptera | native |
| Pilaria | -0.008 | -0.022 | -0.03 | -0.032 | Diptera | native |
| Hexatoma | -0.032 | -0.142 | -0.073 | -0.146 | Diptera | native |
| Ptychoptera | -0.124 | -0.088 | -0.046 | -0.164 | Diptera | native |
| Dixella | 0.095 | -0.103 | -0.081 | -0.033 | Diptera | native |
| Atherix | -0.084 | -0.125 | -0.083 | -0.176 | Diptera | native |
| Atrichops | -0.198 | 0.002 | 0.027 | -0.128 | Diptera | native |
| Limnophora | 0.182 | 0.03 | 0.061 | 0.172 | Diptera | native |
| Sisyra | 0.012 | 0.048 | -0.026 | 0.03 | Neuroptera | native |
| Atyaephyra | 0.091 | 0.153 | 0.008 | 0.17 | Malacostraca | alien |
| Orconectes | -0.022 | 0.05 | 0.054 | 0.039 | Malacostraca | alien |
| Pacifastacus | -0.145 | -0.084 | -0.146 | -0.215 | Malacostraca | alien |
| Echinogammarus | -0.11 | 0.141 | 0.177 | 0.086 | Malacostraca | native |
| Gammarus | -0.021 | -0.181 | 0.143 | -0.079 | Malacostraca | native |
| Niphargus | -0.095 | 0.108 | -0.079 | -0.026 | Malacostraca | native |
| Glossiphonia | -0.062 | -0.021 | -0.021 | -0.066 | Clitellata | native |
| Helobdella | -0.008 | -0.344 | 0.156 | -0.174 | Clitellata | native |
| Piscicola | -0.108 | -0.03 | 0.043 | -0.079 | Clitellata | native |
| Erpobdella | -0.008 | -0.344 | 0.156 | -0.174 | Clitellata | native |
| Theodoxus | -0.133 | 0.232 | -0.082 | 0.028 | Gastropoda | native |
| Valvata | 0.169 | 0.173 | -0.015 | 0.229 | Gastropoda | native |
| Pseudamnicola | 0.103 | 0.129 | -0.008 | 0.156 | Gastropoda | native |
| Potamopyrgus | 0.068 | 0.263 | 0.271 | 0.33 | Gastropoda | alien |
| Belgrandia | -0.018 | 0.155 | -0.054 | 0.07 | Gastropoda | native |
| Emmericia | 0.013 | 0.109 | -0.061 | 0.058 | Gastropoda | alien |
| Bythinella | -0.05 | -0.138 | -0.129 | -0.178 | Gastropoda | native |
| Bithynia | 0.068 | 0.193 | 0.006 | 0.179 | Gastropoda | native |
| Galba | 0.05 | -0.125 | 0.266 | 0.056 | Gastropoda | native |
| Radix | 0.057 | -0.16 | -0.076 | -0.096 | Gastropoda | native |
| Gyraulus | -0.2 | -0.065 | -0.001 | -0.185 | Gastropoda | native |
| Armiger | -0.166 | -0.003 | 0.023 | -0.11 | Gastropoda | native |
| Menetus | -0.203 | 0.034 | 0.019 | -0.113 | Gastropoda | alien |
| Ancylus | -0.155 | 0.026 | 0.09 | -0.057 | Gastropoda | native |
| Ferrissia | 0.08 | 0.139 | 0.198 | 0.227 | Gastropoda | alien |
| Acroloxus | -0.025 | -0.05 | 0.047 | -0.033 | Gastropoda | native |
| Anodonta | -0.092 | -0.075 | 0.053 | -0.094 | Bivalvia | native |
| Unio | 0.018 | -0.011 | 0.045 | 0.023 | Bivalvia | native |
| Pisidium | -0.146 | -0.01 | 0.193 | -0.034 | Bivalvia | native |
| Sphaerium | -0.014 | -0.075 | 0.121 | -0.012 | Bivalvia | native |
| Dreissena | 0.095 | 0.012 | -0.067 | 0.049 | Bivalvia | alien |
| Corbicula | -0.115 | 0.327 | 0.12 | 0.184 | Bivalvia | alien |
| Dugesia | 0.131 | 0.003 | 0.041 | 0.111 | Rhabditophora | native |

| | | | | | | |
|---|---|---|---|---|---|---|
| Polycelis | -0.014 | -0.394 | 0.091 | -0.237 | Rhabditophora | native |
| Dendrocoelum | -0.053 | -0.049 | 0.081 | -0.038 | Rhabditophora | native |
| Hydra | -0.013 | -0.321 | 0.192 | -0.147 | Hydrozoa | native |
| Agriotypus | -0.135 | -0.088 | 0.05 | -0.134 | Hymenoptera | native |
| Procambarus | 0.076 | -0.079 | 0.287 | 0.114 | Malacostraca | alien |
| Torleya | 0.013 | -0.125 | -0.159 | -0.137 | Ephemeroptera | native |
| Hydrobius fuscipes | 0.014 | 0.17 | -0.123 | 0.075 | Coleoptera | native |
| Chalcolestes | 0.063 | 0.077 | -0.044 | 0.079 | Odonata | native |
| Chelifera | -0.206 | -0.065 | -0.017 | -0.196 | Diptera | native |
| Hydrellia | 0.084 | 0.065 | 0.038 | 0.118 | Diptera | native |
| Lispe | -0.074 | -0.015 | 0.088 | -0.028 | Diptera | native |
| Prostoma | -0.008 | 0.272 | -0.032 | 0.163 | Enopla | native |
| Electrogena | -0.009 | -0.022 | -0.101 | -0.061 | Ephemeroptera | native |
| Halesus | -0.124 | -0.088 | -0.046 | -0.164 | Trichoptera | native |
| Raptobaetopus tenellus | -0.116 | 0.069 | 0 | -0.036 | Ephemeroptera | native |
| Pseudocentroptilum | 0.045 | 0.107 | -0.118 | 0.057 | Ephemeroptera | native |
| Corophium | 0.23 | 0.025 | -0.04 | 0.164 | Malacostraca | alien |
| Dikerogammarus | 0.229 | 0.027 | -0.043 | 0.163 | Malacostraca | alien |
| Jaera | -0.027 | -0.051 | 0.079 | -0.022 | Malacostraca | alien |
| Crangonyx | 0.021 | 0.056 | 0.083 | 0.085 | Malacostraca | alien |
| Orchestia | -0.114 | -0.011 | 0.046 | -0.07 | Malacostraca | alien |
| Acentrella | 0.209 | 0.086 | -0.208 | 0.123 | Ephemeroptera | native |
| Serratella | 0.103 | 0.101 | -0.193 | 0.064 | Ephemeroptera | native |
| Scleroprocta | -0.124 | -0.088 | -0.046 | -0.164 | Diptera | native |
| Branchiobdella | -0.117 | -0.069 | 0.001 | -0.128 | Clitellata | native |
| Physella | 0.196 | 0.03 | 0.231 | 0.25 | Gastropoda | alien |
| Limnomysis | 0.166 | 0.004 | 0.002 | 0.121 | Malacostraca | alien |
| Xironogiton | -0.069 | -0.053 | -0.066 | -0.11 | Clitellata | alien |
| Physa stricto-sensu | -0.181 | 0.012 | -0.008 | -0.124 | Gastropoda | native |
| Baetopus | 0.08 | -0.004 | 0.051 | 0.074 | Ephemeroptera | native |
| Pomatinus | 0.088 | 0.066 | -0.086 | 0.072 | Coleoptera | native |

Table S3. Match of trait scores from (p)RDAs with the a priori predictions described Fig. 1.

| driver | model | match | pval |
|---|---|---|---|
| Toxicty | maximal.size | -0.7745 | 0.231 |
| | life.duration | 1.151 | 0.12745 |

| | | | | |
|---|---|---|---|---|
| | cycles.per.year | 0.5495 | 0.301 | |
| | reproduction | 0.206 | 0.412 | |
| | aquatic.stages | 0.244 | 0.3135 | |
| | dispersal | 0.189 | 0.264 | |
| | resistance.forms | -0.693 | 0.2735 | |
| | respiration | 0.4855 | 0.102 | |
| | locomotion | -1.09 | 0.139 | |
| | food | 0.5165 | 0.304 | |
| | feeding.habits | 0.6975 | 0.12 | |
| Temperature | maximal.size | 1.415 | 0.095 | . |
| | life.duration | 0.8155 | 0.231 | |
| | cycles.per.year | 1.91 | 0.027495 | * |
| | reproduction | 0.9 | 0.187 | |
| | aquatic.stages | 0.9465 | 0.1975 | |
| | dispersal | 0.627 | 0.271 | |
| | resistance.forms | 0.773 | 0.2315 | |
| | respiration | 1.64 | 0.0715 | . |
| | locomotion | 1.25 | 0.038 | * |
| | food | 1.52 | 0.0729 | . |
| | feeding.habits | 1.15 | 0.141 | |
| Nutrients | maximal.size | 0.83 | 0.224 | |
| | life.duration | -1.74 | 0.04795 | * |
| | cycles.per.year | 2.44 | 0.004995 | ** |
| | reproduction | 2.67 | 0.000999 | *** |
| | aquatic.stages | 2.85 | 0.000999 | *** |
| | dispersal | 1.15 | 0.149 | |
| | resistance.forms | 1.19 | 0.13095 | |
| | respiration | -0.437 | 0.3485 | |
| | locomotion | 1.11 | 0.147 | |
| | food | -1.56 | 0.05995 | . |
| | feeding.habits | -0.1553 | 0.3855 | |
| Combined | maximal.size | 0.994 | 0.19295 | |
| | life.duration | 0.405 | 0.351 | |
| | cycles.per.year | 2.88 | 0.0015 | ** |
| | reproduction | 1.84 | 0.045 | * |
| | aquatic.stages | 2.145 | 0.02995 | * |
| | dispersal | 1.29 | 0.0999 | . |
| | resistance.forms | 1.107 | 0.143 | |
| | respiration | 1.31 | 0.1025 | |
| | locomotion | 0.925 | 0.139 | |
| | food | 1.14 | 0.129 | |
| | feeding.habits | 0.794 | 0.224 | |
