## Supplementary material for "Uncovering aquatic toxicity as a specific stressor among multiple environmental drivers affecting the functional structure of French stream macroinvertebrate communities": R scripts: Function_var_part.R

```
var.part.threetables <- function(df.resp, list.df.pred, 
                                 provide.var = F,
                                 var.predict1 = colnames(list.df.pred[[1]]), 
                                 var.predict2 = colnames(list.df.pred[[2]]),
                                 var.predict3 = colnames(list.df.pred[[3]]), 
                                 pval.min = rep(0.05, 3),
                                 nperm = 999, max.nb.var = 5){
  
  if(provide.var == T){
    list.df.pred[[1]] <- list.df.pred[[1]][is.element(colnames(list.df.pred[[1]]), var.predict1)]
    list.df.pred[[2]] <- list.df.pred[[2]][is.element(colnames(list.df.pred[[2]]), var.predict2)]
    list.df.pred[[3]] <- list.df.pred[[3]][is.element(colnames(list.df.pred[[3]]), var.predict3)]
    
    list_selected_varnames <- list(var.predict1, var.predict2, var.predict3)
  } else {
    list_selected_varnames <- list()
  }
  
  Rsqrselected_var <- c() # R2 values for each RDA
  Rsqrselected_varAdj <- c() # adjusted R2 values for each RDA
  for (i in 1:7){  # seven RDAs are necessary for partitioning the variance from three predictive datasets
    
    if (i <= 3) {
      # Select one predictive dataset (varpred.df)
      varpred.df <- list.df.pred[[i]]
      selected_varnames <- list_selected_varnames[[i]]
    } else {
      if (i == 4){
        varpred.df <- cbind(list.df.pred[[1]], list.df.pred[[2]]) # tableaux 1 et 2 combinés
        selected_varnames <- c(list_selected_varnames[[1]], list_selected_varnames[[2]])
      } else {
        if (i == 5){
          varpred.df <- cbind(list.df.pred[[1]], list.df.pred[[3]]) # tableaux 1 et 3 combinés
          selected_varnames <- c(list_selected_varnames[[1]], list_selected_varnames[[3]])
        } else {
          if (i == 6){
            varpred.df <- cbind(list.df.pred[[2]], list.df.pred[[3]]) # tableaux 2 et 3 combinés
            selected_varnames <- c(list_selected_varnames[[2]], list_selected_varnames[[3]])
          } else {
            if(i == 7){
              varpred.df <- cbind(list.df.pred[[1]], list.df.pred[[2]], 
                                  list.df.pred[[3]]) # tableaux 1, 2 et 3 combinés
              selected_varnames <- c(list_selected_varnames[[1]], list_selected_varnames[[2]], 
                                     list_selected_varnames[[3]])
            }
          }
        }
      }
    }
    
    if (identical(selected_varnames, character(0))){
      Rsqrselected_var <- c(Rsqrselected_var, 0)
      Rsqrselected_varAdj <- c(Rsqrselected_varAdj, 0)
    } else {
      fmla <- as.formula(paste("df.resp ~ ", paste(selected_varnames, collapse= "+")))
      RDA.resp <- rda(fmla , varpred.df, scale=T) ## RDA

      Rsqrselected_var <- c(Rsqrselected_var, RsquareAdj(RDA.resp)$r.squared)
      Rsqrselected_varAdj <- c(Rsqrselected_varAdj, RsquareAdj(RDA.resp)$adj.r.squared)
    }
  }    
  names(Rsqrselected_varAdj) <- c("ET", "PC", "MEM", "ET+PC", "ET+MEM","PC+MEM","ET+PC+MEM")
  
  ## VARIANCE PARTITIONING
  
  abcd_adj <- Rsqrselected_varAdj["ET"]  #adjR-squ
  ebcf_adj <- Rsqrselected_varAdj["PC"]  #adjR-sqr
  cdfg_adj <- Rsqrselected_varAdj["MEM"]  #adjR-sqr
  
  abcdef_adj <- Rsqrselected_varAdj["ET+PC"]  #adjR-sqr
  abcdfg_adj <- Rsqrselected_varAdj["ET+MEM"] #adjR-sqr
  bcefdg_adj <- Rsqrselected_varAdj["PC+MEM"] #adjR-sqr
  abcdefg_adj <- Rsqrselected_varAdj["ET+PC+MEM"]   #adjR-sqr
  
  abe_adj <- abcdefg_adj - cdfg_adj
  dcf_adj <- abcdef_adj - abe_adj
  
  cd_adj <- abcd_adj + cdfg_adj - abcdfg_adj
  bc_adj <- abcd_adj + ebcf_adj - abcdef_adj
  cf_adj <- ebcf_adj + cdfg_adj - bcefdg_adj
  
  a_adj <- abcdefg_adj - bcefdg_adj
  e_adj <- abcdefg_adj - abcdfg_adj
  g_adj <- abcdefg_adj - abcdef_adj
  c_adj <- abcd_adj + ebcf_adj + cdfg_adj - 2*abcdefg_adj + a_adj + e_adj + g_adj
  b_adj <- bc_adj - c_adj
  d_adj <- cd_adj - c_adj
  f_adj <- cf_adj - c_adj
  
  vecPart <- c(a_adj,b_adj,c_adj,d_adj,e_adj,f_adj,g_adj)
  names(vecPart) <- NULL
  vecPart[vecPart<0] <- 0
  
  return(list(vecPart, list_selected_varnames))
}
```
