## Supplementary material for "Uncovering aquatic toxicity as a specific stressor among multiple environmental drivers affecting the functional structure of French stream macroinvertebrate communities": R scripts: MainRscript_BioRxiv.R

```
library(ade4)
library(spdep)
library(adespatial)
library(vegan)
library(eulerr)
library(FD)
library(infer)
library(rstatix)

## Set directory
setwd("[...]")

## Data available on data.gouv repository : https://doi.org/10.57745/CZTZWJ
DataMIA <- read.csv("MacroinvertebrateSpeciesPerStation.csv", row.names = 1)
DataFIenvir <- read.csv("EnvironmentalDriversPerStation.csv", row.names = 1, col.names = c("", "FI", "Temp","NH4","NO3","NO2","PO4"))
DataEcotox <- DataFIenvir["FI"]
DataEnvir <-  DataFIenvir[c("Temp","NH4","NO3","NO2","PO4")]
TraitsMI <- read.csv("MacroinvertebrateTraitCategoryPerSpecies.csv", row.names = 1)
rownames(TraitsMI) <- gsub("[ -]", ".", rownames(TraitsMI))
CWMtraitsMI <- read.csv("MacroinvertebrateTraitCWMPerStation.csv", row.names = 1)
TaxoMIA <- read.csv("MacroinvertebrateTaxonomy.csv", row.names = 1)


## Script on BioRxiv : https://doi.org/10.64898/2026.01.08.698385
## Load variation partitioning function 
source("[...]/Function_var_part.R")

# Hellinger transformation
# (for each station, abundances divided by station total abundance, then square root transformation)
L.MIAHell <- decostand(DataMIA, method = "hellinger")

# z-scores 
# (each variable as mean = 0 and sd = 1)
R.Ecotox.zsco <- decostand(log(DataEcotox), method = "standardize") 

R.Envir.zsco <- decostand(log(DataEnvir), method = "standardize") 
 
## PCA on nutrient variables 
Nutrients.PCA <- rda(R.Envir.zsco[c("NH4", "NO3", "NO2", "PO4")])
plot(Nutrients.PCA, choices = c(1,2))
# Creation of the "Nutrients" variable from the PCA on the "NH4", "NO3", "NO2" and "PO4" variables
Nutrients.PCA1 <- scores(Nutrients.PCA, choices = c(1), display = c("wa"),scaling = "species")
colnames(Nutrients.PCA1) <- "Nutrients"

## PCA on all drivers
drivers.PCA <- rda(cbind(R.Ecotox.zsco, R.Envir.zsco["Temp"], Nutrients.PCA1))
plot(drivers.PCA, choices = c(1, 2))
plot(drivers.PCA, choices = c(2, 3))
# Creation of the "drivers" variable from the PCA on all variables
drivers.PCA2 <- scores(drivers.PCA, choices = c(2), display = c("wa"),scaling = "species")
colnames(drivers.PCA2) <- "drivers"


############################################################################################
## Variation partitioning on macroinvertebrate taxonomical and functional data

var.drivers <- c("FI", "Temp", "Nutrients")
df.resp_2 <- list(L.MIAHell, CWMtraitsMI)
names(df.resp_2) <- c("taxo", "traits")
list.df.pred_2 <- list(R.Ecotox.zsco, R.Envir.zsco["Temp"], as.data.frame(Nutrients.PCA1))

all.frac <- list()
list.varparts <- list()
list.results <- list()
for (j in 1:length(df.resp_2)){
  ##################################################################################################
  result.var.part <- var.part.threetables(df.resp = df.resp_2[[j]], 
                                          list.df.pred = list.df.pred_2,
                                          provide.var = T, 
                                          var.predict1 = var.drivers[1],   # "FI"
                                          var.predict2 =  var.drivers[2],   # "Temp" 
                                          var.predict3 = var.drivers[3],  # "Nutrients" 
                                          nperm = 999)
  ##################################################################################################
  result.var.part ## Fractions of explained variance as adjusted Rsqr values
  
  list.varparts <- c(list.varparts, list(result.var.part))
  all.frac <- c(all.frac, list(c(sum(result.var.part[[1]][1:4]), 
                            sum(result.var.part[[1]][c(2,3,5,6)]),
                            sum(result.var.part[[1]][c(3,4,6,7)])
                            )))

  list.sco.pur <- list()       ## list of species or trait scores from pRDA (pure fraction of explained variance only)
  list.sco.sta.pur <- list()   ## list of station scores from pRDA (pure fraction of explained variance only)
  list.scores <- list()        ## list of species or trait scores from RDA (explained variance including shared fractions)
  list.sco.stations <- list()  ## list of station scores from RDA (explained variance including shared fractions)
  list.rda.pur    <- list()    ## list of pRDA results (pure fraction of explained variance only)
  list.rda    <- list()        ## list of RDA results (explained variance including shared fractions)
  purefracanova <- list()      ## Results of permutation tests on the pRDA (pure fraction of explained variance only)
  allfracanova <- list()       ## Results of permutation tests on the RDA (explained variance including shared fractions)
  for (i in 1:length(var.drivers)){
    
    ##################################################################
    ## Significativity of the variance explained variance by each variable independently from each other
    ###
    # partial RDA with one variable as predictor and the two others as conditional variables
    nom.var.drivers.pred <- var.drivers[i] # predictor variable
    nom.var.drivers.cond <- var.drivers[-i] # conditional variables
    varpred.df <- cbind.data.frame(list.df.pred_2[[i]], list.df.pred_2[-i])
    fmla <- as.formula(paste0(expression(df.resp_2[[j]])," ~ ", paste(nom.var.drivers.pred, collapse= " + "),
                              " + Condition(", paste(nom.var.drivers.cond, collapse= ") + Condition("), ")"))
    RDA.MIA_pure <- rda(fmla , varpred.df, scale=T) ## pRDA
    perm = how(nperm=999) ## 
    anovaRDA.MIA_pure <- anova(RDA.MIA_pure, permutations=perm) # Permutation test
    
    ##################################################################
    ## Significativity of the variance explained by each variable, including shared variance between variables
    fmla2 <- as.formula(paste0(expression(df.resp_2[[j]])," ~ ", paste(nom.var.drivers.pred, collapse= " + ")))
    RDA.MIA <- rda(fmla2 , varpred.df, scale=T) ## RDA
    anovaRDA.MIA <- anova(RDA.MIA, permutations=perm) # Permutation test
    
    ## 
  
    purefracanova <- c(purefracanova, list(anovaRDA.MIA_pure))
    allfracanova <- c(allfracanova, list(anovaRDA.MIA))
    
    scores.RDA.pur <- scores(RDA.MIA_pure, display = c("sp"), scaling = "species")
    scores.sta.pur <- scores(RDA.MIA_pure, display = c("sites"))
    
    scores.RDA.MIA <- scores(RDA.MIA, display = c("sp"), scaling = "species")
    scores.stations <- scores(RDA.MIA, display = c("sites"))
    
    if (RDA.MIA_pure$CCA$biplot[,1] < 0){
      RDA.MIA_pure$CCA$biplot[,1] <- -RDA.MIA_pure$CCA$biplot[,1]
      scores.RDA.pur[,1] <- -scores.RDA.pur[,1]
      scores.sta.pur[,1] <- -scores.sta.pur[,1]
    }
    
    if (RDA.MIA$CCA$biplot[,1] < 0){
      RDA.MIA$CCA$biplot[,1] <- -RDA.MIA$CCA$biplot[,1]
      scores.RDA.MIA[,1] <- -scores.RDA.MIA[,1]
      scores.stations[,1] <- -scores.stations[,1]
    }
    
    list.sco.pur <- c(list.sco.pur, list(scores.RDA.pur))
    list.sco.sta.pur <- c(list.sco.sta.pur, list(scores.sta.pur))
    list.scores <- c(list.scores, list(scores.RDA.MIA))
    list.sco.stations <- c(list.sco.stations, list(scores.stations))
    list.rda.pur  <- c(list.rda.pur, list(RDA.MIA_pure))
    list.rda    <- c(list.rda, list(RDA.MIA))
    
    names(list.sco.pur)[i] <- var.drivers[i]
    names(list.sco.sta.pur)[i] <- var.drivers[i]
    names(list.rda.pur)[i]    <- var.drivers[i]
    names(list.scores)[i] <- var.drivers[i]
    names(list.sco.stations)[i] <- var.drivers[i]
    names(list.rda)[i]    <- var.drivers[i]
    names(purefracanova)[i] <- var.drivers[i]
    names(allfracanova)[i] <- var.drivers[i]
  }
  
  list.results[[j]] <- list(list.sco.pur, list.sco.sta.pur, list.rda.pur, 
                                    list.scores, list.sco.stations, list.rda, 
                                    purefracanova, allfracanova)
  names(list.results[[j]]) <- c("sco.pur", "sco.sta.pur", "rda.pur",
                                        "scores", "sco.stations", "rda", 
                                        "purefracanova", "allfracanova")
}
names(all.frac) <- names(df.resp_2)
names(list.varparts) <- names(df.resp_2)
names(list.results) <- names(df.resp_2)


#######################################################################
## Combined effects of all drivers on on macroinvertebrates

df.resp_2 <- list(L.MIAHell, CWMtraitsMI)
names(df.resp_2) <- c("taxo", "traits")
list.scores.all <- list()
list.sco.stations.all <- list()
list.rda.all    <- list()
alldriversanova <- list()
for (j in 1:length(df.resp_2)){
  nom.var.drivers.pred <- "drivers"
  varpred.df <- as.data.frame(drivers.PCA2)
  fmla2 <- as.formula(paste0(expression(df.resp_2[[j]])," ~ ", paste(nom.var.drivers.pred, collapse= "+")))
  RDA.MIA <- rda(fmla2 , varpred.df, scale=T) ## RDA
  
  perm = how(nperm=999) ## 
  anovaRDA.MIA <- anova(RDA.MIA, permutations=perm) #Permutation test

  scores.RDA.MIA <- scores(RDA.MIA, display = c("sp"), scaling = "species", choices = c(1,2,3))
  scores.stations <- scores(RDA.MIA, display = c("sites"), choices = c(1,2,3))

  if (mean(RDA.MIA$CCA$biplot) < 0){
    RDA.MIA$CCA$biplot <- -RDA.MIA$CCA$biplot
    scores.RDA.MIA <- -scores.RDA.MIA
    scores.stations <- -scores.stations
  }

  list.scores.all <- c(list.scores.all, list(scores.RDA.MIA))
  names(list.scores.all)[j] <- names(df.resp_2)[j]

  list.sco.stations.all <- c(list.sco.stations.all, list(scores.stations))
  names(list.sco.stations.all)[j] <- names(df.resp_2)[j]

  list.rda.all    <- c(list.rda.all, list(RDA.MIA))
  names(list.rda.all)[j] <- names(df.resp_2)[j]
  
  alldriversanova <- c(alldriversanova, list(anovaRDA.MIA))
  names(alldriversanova)[j] <- names(df.resp_2)[j]
}


#################################################################
## Summary results variation partitioning

for (i in 1:length(list.varparts)){
  if (i == 1){
    tab.varpart <- list.varparts[[i]][[1]]
  } else {
    tab.varpart <- rbind(tab.varpart, list.varparts[[i]][[1]])
  }
}
rownames(tab.varpart) <- c("Taxo", "Traits")
colnames(tab.varpart) <- c("FI", "FI+Temp.", "FI+Temp.+Nutrient", "FI+Nutrient", "Temp.", "Temp.+Nutrient", "Nutrient")

totvar <- sapply(as.data.frame(t(tab.varpart)), sum)
EToutENV <- (tab.varpart[,1] + tab.varpart[,4])/totvar*100
ETwithENV <- (tab.varpart[,2] + tab.varpart[,3])/totvar*100
ENVoutET <- (tab.varpart[,5] + tab.varpart[,6])/totvar*100

EToutMEM <- (tab.varpart[,1] + tab.varpart[,2])/totvar*100
ETwithMEM <- (tab.varpart[,3] + tab.varpart[,4])/totvar*100
MEMoutET <- (tab.varpart[,6] + tab.varpart[,7])/totvar*100

EToutENVMEM <- (tab.varpart[,1])/totvar*100
ETwithENVoutMEM <- (tab.varpart[,2])/totvar*100
ETwithENVMEMout <- (tab.varpart[,3])/totvar*100
ETwithMEMoutENV <- (tab.varpart[,4])/totvar*100
ETwithENVMEM <- (tab.varpart[,2] + tab.varpart[,3] + tab.varpart[,4])/totvar*100
ENVoutETMEM <- (tab.varpart[,5])/totvar*100
ENVwithMEMoutET <- (tab.varpart[,6])/totvar*100
MEMoutETENV <- (tab.varpart[,7])/totvar*100

Relativ.Part1 <- rbind(EToutENV, ETwithENV, ENVoutET)
Relativ.Part1 <- Relativ.Part1[,ncol(Relativ.Part1):1]
Relativ.Part2 <- rbind(EToutMEM, ETwithMEM, MEMoutET)
Relativ.Part2 <- Relativ.Part2[,ncol(Relativ.Part2):1]

Relativ.Part3 <- rbind(EToutENVMEM, ETwithENVoutMEM, ETwithMEMoutENV, ETwithENVMEMout, ENVoutETMEM, ENVwithMEMoutET, MEMoutETENV)
Relativ.Part3 <- Relativ.Part3[,ncol(Relativ.Part3):1]

par(mar=c(4, 4, 2, 10) + 0.1)
bp <- barplot(Relativ.Part3, horiz = T, axisnames = F, axes=T, 
              col = c(rgb(255,153,153, maxColorValue = 255),
                      rgb(255,173,173, maxColorValue = 255),
                      rgb(255,173,163, maxColorValue = 255),
                      rgb(255,173,153, maxColorValue = 255),
                      rgb(255,255,153, maxColorValue = 255),
                      rgb(233,233,153, maxColorValue = 255),
                      rgb(153,153,255, maxColorValue = 255)),
              space = c(0.5), 
              xlim = c(0, 100))

text(x = -1, y = bp, labels = colnames(Relativ.Part2), cex = 1, adj = c(1, 0.5), xpd = T)
mtext("% of the total explained variance", side = 1, line = 2.5)


for (j in 1:ncol(Relativ.Part3)){
  cumulx <- Relativ.Part3[1,j]
  for (i in 2:nrow(Relativ.Part3)){
    cumulx <- c(cumulx, sum(Relativ.Part3[1:(i-1),j]) + Relativ.Part3[i,j])
  }
  text(x = cumulx[Relativ.Part3[,j] > 1.2] - 0.7, y = bp[j]-0.3, cex = 0.6,
       labels = which(Relativ.Part3[,j] > 1.2))
}


legend(x = 105, y = bp[bp==max(bp)]+0.5, c("(1) FI",
                                           "(2) FI+Temp.",
                                           "(3) FI+Nutrient",
                                           "(4) FI+Temp.+Nutrients",
                                           "(5) Temp.", 
                                           "(6) Temp.+Nutrients",
                                           "(7) Nutrients"
), 
fill = c(rgb(255,153,153, maxColorValue = 255),
         rgb(255,173,173, maxColorValue = 255),
         rgb(255,173,163, maxColorValue = 255),
         rgb(255,173,153, maxColorValue = 255),
         rgb(255,255,153, maxColorValue = 255),
         rgb(233,233,153, maxColorValue = 255),
         rgb(153,153,255, maxColorValue = 255)
),
title = "Fractions:", title.adj = 0, title.cex = 1, cex = 0.9, title.col = "black", xjust = 0, bty ="n", xpd=T)


## Taxon scores RDA Taxo
TaxscoRDAtaxo_FI <- list.results$taxo$sco.pur$FI[,1]
TaxscoRDAtaxo_Temp <- list.results$taxo$sco.pur$Temp[,1]
TaxscoRDAtaxo_Nutrient <- list.results$taxo$sco.pur$Nutrient[,1]
TaxscoRDAtaxo_ALL <- list.scores.all$taxo[,1]
## Traits scores RDA Traits
TrascoRDAtraits_FI <- list.results$traits$sco.pur$FI[,1]
TrascoRDAtraits_Temp <- list.results$traits$sco.pur$Temp[,1]
TrascoRDAtraits_Nutrient <- list.results$traits$sco.pur$Nutrient[,1]
TrascoRDAtraits_ALL <- list.scores.all$traits[,1]

## Pearson's correlations between scores
cor.test(TaxscoRDAtaxo_ALL, TaxscoRDAtaxo_FI)
cor.test(TaxscoRDAtaxo_ALL, TaxscoRDAtaxo_Temp)
cor.test(TaxscoRDAtaxo_ALL, TaxscoRDAtaxo_Nutrient)
cor.test(TrascoRDAtraits_ALL, TrascoRDAtraits_FI)
cor.test(TrascoRDAtraits_ALL, TrascoRDAtraits_Temp)
cor.test(TrascoRDAtraits_ALL, TrascoRDAtraits_Nutrient)


####################################################################################
## Comparison with a priori and null models

Model_apriori        <- vector(mode = "numeric", length = ncol(TraitsMI))
names(Model_apriori) <- colnames(TraitsMI)
## Size
Model_apriori[c("inf.0.25.cm", "sup.0.25.0.5.cm", "sup.0.5.1.cm", "sup.1.2.cm")] <- 
  c(1            , 1                , -1            , -1           )
## Life
Model_apriori[c("inf.1.year", "sup.1.year")] <- 
  c( 1          , -1          )
## Cycle
Model_apriori[c("inf.1.cy", "sup.1.cy")] <- 
  c( -1        , 1        )
## Repro
Model_apriori[c("ovo", "tec", "cie")] <- 
  c(1    , -1   , -1   )
## Aqua
Model_apriori[c("larva", "adult")] <- 
  c(-1     , 1      )
## Disper
Model_apriori[c("aquatic.passive", "aerial.active", "aquatic.active")] <- 
  c(1                , -1             , -1              )
## Resis
Model_apriori[c("egs", "non", "hou", "did")] <- 
  c(-1    ,-1   , 1    , 1    )
## Respi
Model_apriori[c("gil", "spi")] <- 
  c(1    , - 1  )
## Loco
Model_apriori[c("crw", "int", "bur")] <- 
  c(-1    , 1   , 1    )
## Food
Model_apriori[c("lmac", "vert", "detl1", "mior")] <- 
  c(-1    , -1    , 1      , 1     )
## Feeding
Model_apriori[c("pre", "par", "fif", "dpf")] <- 
  c(-1   , -1   , 1    , 1    )

Model_apriori <- Model_apriori[Model_apriori != 0]

################################################################################


## Function which create null trait matrix
shuffletraitvalues <- function(TraitMatrix, model = 1){
  shuffludTraitMatrix <- TraitMatrix
  if (model == 1){
    shuffludTraitMatrix <- shuffludTraitMatrix[sample(nrow(TraitMatrix)), ]
    rownames(shuffludTraitMatrix) <- rownames(TraitMatrix)
  } else {
    if (model == 2){
      for (i in 1:ncol(TraitMatrix)){
        shuffludTraitMatrix[,i] <- sample(TraitMatrix[,i])
      }
    }
  }
  return(shuffludTraitMatrix)
}

## Probability station x abundance matrix
mgen <- function(m, n, e = TRUE, zs = FALSE){   # zs = T allows species with no interaction (empty rows or columns)
  mac <- matrix(cumsum(m), nrow(m), ncol(m)) #Cumulative probability matrix
  mint <- matrix(0, nrow(m), ncol(m)) #Interaction matrix
  
  if (zs == FALSE){
    for (i in 1:nrow(m)){
      c1<-sample(ncol(m),replace=T,prob=colSums(m)); c1<-c1[1]
      mint[i,c1]<-1
    }
    for (i in 1:ncol(m)){
      r1<-sample(nrow(m),replace=T,prob=rowSums(m)); r1<-r1[1]
      mint[r1,i]<-1
    }
  }
  while(sum(mint)<n){
    rand<-runif(1,0,1)
    ri<-min(which(mac>=rand))
    mint[ri]<-mint[ri]+1
  }
  return(mint)
}

###########################################################################################################
## Create null CWM matrices and do RDAs on them with the different drivers alone or in combination
Nrep <- 100 #1000 Can be long
var.drivers <- c("FI", "Temp", "Nutrients", "ALL")
list.sco.traits <- list(list(), list(), list(), list())
names(list.sco.traits) <- var.drivers
listAdjR2 <- list()
for (k in 0:Nrep){
  if (k == 0){
    df.resp <- CWMtraitsMI
  } else {
    MIA.new <- mgen(m = as.matrix(DataMIA+1)/sum(DataMIA), n = 10000, e = TRUE, zs = F)
    rownames(MIA.new) <- rownames(DataMIA)
    colnames(MIA.new) <- colnames(DataMIA)
    MIA.newHell <- decostand(MIA.new, method = "hellinger")
    
    TraitsMI_RANDOM <- shuffletraitvalues(TraitsMI, model = 1)
    NamesBinaryTraits <- colnames(TraitsMI_RANDOM[sapply(TraitsMI_RANDOM, max)==1])
    rownames(TraitsMI_RANDOM) 
    CWMtraitsMI_RANDOM <- functcomp(x = TraitsMI_RANDOM, a = as.matrix(MIA.newHell),
                                    CWM.type = "all", bin.num = NamesBinaryTraits)
    CWMtraitsMI_RANDOM[is.na(CWMtraitsMI_RANDOM)] <- 0
    
    df.resp <- CWMtraitsMI_RANDOM
  }
  
  for (var.select in 1:length(var.drivers)){
    if (var.select <= 3){
      var.1 = var.drivers[var.select] 
      var.23 = var.drivers[-var.select]
      var.2 = var.23[1]
      var.3 = var.23[2]
      
      varpred.df <- cbind(R.Ecotox.zsco, R.Envir.zsco["Temp"], as.data.frame(Nutrients.PCA1)) 
      fmla <- as.formula(paste0("df.resp ~ ", paste(var.1, collapse= " + "),
                                " + Condition(", paste(c(var.2, var.3), 
                                                       collapse= ") + Condition("), ")"))
      fmla
      RDA.MIA <- rda(fmla , varpred.df, scale = T) ## RDA
      RDA.MIA
      
      
      ##################################################################
      
      scores.traits <- scores(RDA.MIA, display = c("sp"))
      scores.drivers <- scores(RDA.MIA, display = c("bp"))
      if (scores.drivers[,"RDA1"] < 0){
        scores.traits <- -scores.traits
        scores.drivers <- -scores.drivers
      }
      
    } else {
      #######################################################################
      ## Combined effects of all drivers on macroinvertebrates
      varpred.df <- as.data.frame(drivers.PCA2)
      nom.var.pred <- "drivers"
      fmla2 <- as.formula(paste0("df.resp ~ ", paste(nom.var.pred, collapse= "+")))
      fmla2
      RDA.MIA <- rda(fmla2 , varpred.df, scale=T) ## RDA
      RDA.MIA
      
      scores.traits <- scores(RDA.MIA, display = c("sp"))
      scores.drivers <- scores(RDA.MIA, display = c("bp"))
      if (scores.drivers[,"RDA1"] < 0){
        scores.traits <- -scores.traits
        scores.drivers <- -scores.drivers
      }
      
      ##################################################################
    }
    
    adjR2.RDA <- RsquareAdj(RDA.MIA)$adj.r.squared
    if (k == 0){
      listAdjR2[[var.select]] <- adjR2.RDA
    } else {
      listAdjR2[[var.select]] <- c(listAdjR2[[var.select]], adjR2.RDA)
    }
    
    list.sco.traits[[var.drivers[var.select]]][[k+1]] <- scores.traits
  }
}


######################################################
## MEAN + SD OF NULL MODELS FOR EACH TRAIT CATEGORY
list.table.traits <- list()
for (var.select in 1:length(list.sco.traits)){
  table.traits           <- data.frame(matrix(nrow = ncol(CWMtraitsMI), ncol = 5))
  colnames(table.traits) <- c("obs", "null(mean)", "null(sd)", "std", "pval")
  rownames(table.traits) <- colnames(CWMtraitsMI)
  for (i in 1:ncol(CWMtraitsMI)){
    traitscores <- c()
    for (j in 2:(Nrep+1)){
      traitscores <- c(traitscores, list.sco.traits[[var.drivers[var.select]]][[j]][i,"RDA1"])
    }
    
    trait.name <- rownames(list.sco.traits[[var.drivers[var.select]]][[j]])[i]
    trait.name
    obs.trait <- list.sco.traits[[var.drivers[var.select]]][[1]][i,"RDA1"]
    obs.trait
    mean.null <- mean(traitscores) 
    mean.null
    sd.null   <- sd(traitscores)
    sd.null
    sco.std <- (obs.trait - mean.null)/sd.null
    sco.std 
    
    pval.greater <- 1-sum(c(obs.trait - traitscores)>0)/(Nrep+1) #p value CWM-sp
    pval.greater
    pval.less <- sum(c(obs.trait - traitscores)>0)/(Nrep+1) #p value CWM-sp
    pval.less
    if (pval.greater < pval.less){
      pval.choice <- pval.greater
    } else {
      pval.choice <- pval.less
    }
    pval.choice
    
    table.traits[trait.name, 1:5] <- signif(c(obs.trait, mean.null, sd.null, sco.std, pval.choice), 3)
  }
  
  list.table.traits[[var.select]] <- table.traits
}
names(list.table.traits) <- var.drivers
list.table.traits


######################################################################
## MATCH OF OBSERVED TRAITS WITH A PRIORI
table.match <- data.frame(matrix(nrow = length(list.table.traits)*length(Model_apriori), ncol = 4))
colnames(table.match) <- c("driver", "trait", "match", "pval")
i <- 1
for (var.select in 1:length(list.table.traits)){
  for (model.selec in 1:length(Model_apriori)){
    table.match$driver[[i]] <- var.drivers[var.select]
    table.match$trait[[i]]  <- names(Model_apriori)[model.selec]
    table.match$match[[i]] <- median(list.table.traits[[var.select]][names(Model_apriori)[model.selec], "std"]*Model_apriori[[model.selec]])
    table.match$pval[[i]] <- median(list.table.traits[[var.select]][names(Model_apriori)[model.selec], "pval"])
    i <- i+1
  }
}
table.match


###########################################################################################
## Association between driver-related taxon turnover and diversity metrics

    ## Taxonomic richness (number of genus)
TotSpRichStations <- specnumber(DataMIA)
    ## Taxonomic distances
TaxoDist <- taxa2dist(TaxoMIA, varstep = T, check = TRUE)
    ## Taxonomic distincness indices
TaxoDiv <- taxondive(DataMIA, TaxoDist, match.force = F)
    ## Functional diversity indices
resFD <- dbFD(x = TraitsMI, a = as.matrix(L.MIAHell), calc.CWM = F, stand.FRic = T)

## Station scores RDA Taxo
StascoRDAtaxo_FI <- list.results$taxo$sco.sta.pur$FI[,1]
StascoRDAtaxo_Temp <- list.results$taxo$sco.sta.pur$Temp[,1]
StascoRDAtaxo_Nutrient <- list.results$taxo$sco.sta.pur$Nutrient[,1]
StascoRDAtaxo_ALL <- list.sco.stations.all$taxo[,1]

## Remove outliers in station scores
for (i in 1:2){
  if(any(is_outlier(StascoRDAtaxo_FI))){
    StascoRDAtaxo_FI <- StascoRDAtaxo_FI[!is_outlier(StascoRDAtaxo_FI)]
  }
  if(any(is_outlier(StascoRDAtaxo_Temp))){
    StascoRDAtaxo_Temp <- StascoRDAtaxo_Temp[!is_outlier(StascoRDAtaxo_Temp)]
  }
  if(any(is_outlier(StascoRDAtaxo_Nutrient))){
    StascoRDAtaxo_Nutrient <- StascoRDAtaxo_Nutrient[!is_outlier(StascoRDAtaxo_Nutrient)]
  }
  if(any(is_outlier(StascoRDAtaxo_ALL))){
    StascoRDAtaxo_ALL <- StascoRDAtaxo_ALL[!is_outlier(StascoRDAtaxo_ALL)]
  }
}

    ## Linear models
    ## Taxonomic richness ~ station scores
summary(lm(TotSpRichStations[names(StascoRDAtaxo_FI)] ~ StascoRDAtaxo_FI))
summary(lm(TotSpRichStations[names(StascoRDAtaxo_Temp)] ~ StascoRDAtaxo_Temp))
summary(lm(TotSpRichStations[names(StascoRDAtaxo_Nutrient)] ~ StascoRDAtaxo_Nutrient))
summary(lm(TotSpRichStations[names(StascoRDAtaxo_ALL)] ~ StascoRDAtaxo_ALL))
    ## Taxonomic distinctness ~ station scores
summary(lm(TaxoDiv$Dstar[names(StascoRDAtaxo_FI)] ~ StascoRDAtaxo_FI))
summary(lm(TaxoDiv$Dstar[names(StascoRDAtaxo_Temp)] ~ StascoRDAtaxo_Temp))
summary(lm(TaxoDiv$Dstar[names(StascoRDAtaxo_Nutrient)] ~ StascoRDAtaxo_Nutrient))
summary(lm(TaxoDiv$Dstar[names(StascoRDAtaxo_ALL)] ~ StascoRDAtaxo_ALL))
    ## Functional richness ~ station scores
summary(lm(resFD$FRic[names(StascoRDAtaxo_FI)] ~ StascoRDAtaxo_FI))
summary(lm(resFD$FRic[names(StascoRDAtaxo_Temp)] ~ StascoRDAtaxo_Temp))
summary(lm(resFD$FRic[names(StascoRDAtaxo_Nutrient)] ~ StascoRDAtaxo_Nutrient))
summary(lm(resFD$FRic[names(StascoRDAtaxo_ALL)] ~ StascoRDAtaxo_ALL))
    ## Functional dispersion ~ station scores
summary(lm(resFD$FDis[names(StascoRDAtaxo_FI)] ~ StascoRDAtaxo_FI))
summary(lm(resFD$FDis[names(StascoRDAtaxo_Temp)] ~ StascoRDAtaxo_Temp))
summary(lm(resFD$FDis[names(StascoRDAtaxo_Nutrient)] ~ StascoRDAtaxo_Nutrient))
summary(lm(resFD$FDis[names(StascoRDAtaxo_ALL)] ~ StascoRDAtaxo_ALL))
```
